# *Grb7*, *Grb10* and *Grb14,* encoding the growth factor receptor-bound 7 family of signalling adaptor proteins have overlapping functions in the regulation of fetal growth and post-natal glucose metabolism

**DOI:** 10.1101/2024.01.31.578179

**Authors:** Kim Moorwood, Florentia M. Smith, Alastair S. Garfield, Michael Cowley, Lowenna J. Holt, Roger J. Daly, Andrew Ward

## Abstract

**Background:** The growth factor receptor bound 7 (Grb7) family of signalling adaptor proteins comprises Grb7, Grb10 and Grb14. Each can interact with the insulin receptor and other receptor tyrosine kinases, where Grb10 and Grb14 inhibit insulin receptor activity. In cell culture studies they mediate functions including cell survival, proliferation, and migration. Mouse knockout (KO) studies have revealed physiological roles for *Grb10* and *Grb14* in glucose-regulated energy homeostasis. Both *Grb10* KO and *Grb14* KO mice exhibit increased insulin signalling in peripheral tissues, with increased glucose and insulin sensitivity and a modestly increased ability to clear a glucose load. In addition, *Grb10* strongly inhibits fetal growth such that at birth *Grb10* KO mice are 30% larger by weight than wild type littermates.

**Results:** Here, we generate a *Grb7* KO mouse model. We show that during fetal development the expression patterns of Grb7 and Grb14 each overlap with that of Grb10. Despite this, *Grb7* and *Grb14* did not have a major role in influencing fetal growth, either alone or in combination with *Grb10*. At birth, in most respects both *Grb7* KO and *Grb14* KO single mutants were indistinguishable from wild type, while *Grb7*:*Grb10* double knockout (DKO) were near identical to *Grb10* KO single mutant and *Grb10*:*Grb14* DKO mutants slightly smaller. In the developing kidney *Grb7* had a subtle positive influence on growth. An initial characterisation of *Grb7* KO adult mice revealed sexually dimorphic effects on energy homeostasis, with females having significantly smaller white adipose tissue (WAT) depots and an enhanced ability to clear glucose from the circulation, compared to wild type littermates. Males had elevated fasted glucose levels with a trend towards smaller WAT depots, without improved glucose clearance.

**Conclusions:** *Grb7* and *Grb14* do not have significant roles as inhibitors of fetal growth, unlike *Grb10*, and instead *Grb7* may promote growth of the developing kidney. In adulthood, *Grb7* contributes subtly to glucose mediated energy homeostasis, raising the possibility of redundancy between all three adaptors in physiological regulation of insulin signalling and glucose handling.

## Introduction

The growth factor receptor bound (Grb) 7-family comprises Grb7, Grb10 and Grb14, a structurally related group of signalling adaptor proteins [1, 2]. Grb7-family proteins lack catalytic activity but share several conserved molecular interaction domains, that are similarly ordered from amino– (N-) to carboxyl– (C-) terminus. These include a proline-rich region towards the N-terminus containing SH3 binding motifs that may be functional [3] and tandem GYF motifs that were found to bind two proteins dubbed Grb10-interacting GYF (GIGYF) binding proteins, GIGYF1 and GIGYF2 [4]. Moving towards the C-terminus, there is a Ras-association-(RA-) like domain that mediates interactions with various signalling molecules, including different Ras proteins (e.g. [5, 6]). Next there is a pleckstrin homology (PH) domain that provides a means of associating with specific membrane inositol phospholipids, that appears to be important for membrane localization and recruitment to the insulin receptor (Insr) [6, 7]. The PH domain is also required for binding to non-receptor intracellular signalling proteins including N-Ras [6] and calmodulin, the latter in a calcium-dependent manner [8, 9]. Nearest the C-terminus is a Src-homology 2 (SH2) domain that mediates interactions with activated receptor tyrosine kinases (RTKs) and other signalling proteins [1]. Finally, a characteristic feature of the Grb7-family is a conserved region not found in other proteins that lies between the PH and SH2 domains, termed the BPS domain, that also plays a role in RTK interactions, with the N-terminal portion occupying the Insr kinase substrate groove and thereby acting as a pseudosubstrate inhibitor [10].

Despite the SH2 domain being the most highly conserved region between the three Grb7-family members they each exhibit preferential binding to an overlapping set of RTKs, though it should be noted that a variety of approaches have been used and few studies directly compare binding of two or more members to the same receptor [1]. The most widely studied interactions have been those with the Insr, which all three Grb7-family proteins are capable of binding, and the closely related insulin-like growth factor 1 receptor (Igf1r), mainly studied in relation to Grb10 (reviewed in [11]). Grb10 and Grb14 have emerged as inhibitors of these receptors, influencing downstream signalling pathway molecules that mediate energy metabolism, such as Irs1, p85-PI3K and Akt, as well as those for cell survival and proliferation, notably ERK/MAPK [1]. In addition, Grb10 is a direct substrate for phosphorylation by the mTORC1 growth factor and nutrient sensing complex [12, 13].

Physiological roles for *Grb10* and *Grb14* have been identified through mouse knockout (KO) studies. *Grb10* is one of around 150 mouse genes subject to regulation by genomic imprinting such that usually only one of the two parental alleles is expressed [14, 15]. *Grb10* is unusual in being widely expressed from the maternal allele in developing mesodermal and endodermal tissues, whereas the paternal allele is expressed in the developing central nervous system [16–18]. Consequently, mice inheriting a paternal *Grb10* null allele (*Grb10^+/p^* KO mice) exhibit changes in specific behavioural traits [18–21], whereas those inheriting a mutant maternal allele (*Grb10^m/+^*, hereafter *Grb10* KO) are characterised by fetal and placental overgrowth, such that they are around 30% heavier at birth than wild type littermates [17, 18, 22, 23]. This involves enlargement of endodermal and mesodermal organs but not the brain, consistent with the expression pattern of the maternal *Grb10* allele. The overgrowth of *Grb10* KO mice involves increased cell proliferation and cell number [22, 24] and at birth they have larger skeletal muscles with more myofibers, rather than bigger myofibers, with unaltered ratios of fast and slow-twitch fibres [25]. *Grb10* KO mice retain an increased lean mass profile as adults and have a modest improvement in glucose and insulin sensitivity that is associated with elevated glucose-stimulated Insr signalling [22, 26, 27]. Further, tissue-specific knockouts have shown that disruption of the maternal *Grb10* allele in adipose tissue (brown and white together), pancreas, skeletal muscle or hypothalamus is sufficient to alter different aspects of energy homeostasis [28–31].

Growth regulation by Grb10 during fetal development is widely assumed to occur through inhibition of Igf1r, which is known to act as a major regulator of fetal growth by mediating the growth promoting effects of the two insulin-like growth factor ligands Igf1 and Igf2 (reviewed in [32]). However, tests for epistatic interactions between Grb10 and IGF signaling components, including between *Grb10* KO and either *Igf2* KO [17] or *Igf1r* KO [23] mice provide strong evidence that Grb10 regulates fetal growth largely independently of IGF signaling. In *Grb10* KO mice, the neonatal liver is disproportionately enlarged, with hepatocytes filled with lipid [22]. These hepatic phenotypes were abrogated in *Grb10*:*Insr* DKO mice, indicating that Grb10 normally restricts hepatic lipid storage by inhibiting the Insr, but there was no evidence for involvement of the Insr in other aspects of fetal growth [23].

*Grb14^-/-^* homozygous (*Grb14* KO) mutant mice have a normal lean to adipose body composition and also exhibit modestly improved glucose and insulin sensitivity, with increased glucose-regulated insulin receptor signalling [33, 34]. Adult *Grb10*:*Grb14* double knockout (DKO) mice have the altered body composition characteristic of *Grb10* KO mice, with glucose and insulin sensitivity further improved in comparison to either *Grb10* KO or *Grb14* KO single mutants, suggesting an additive effect of losing insulin receptor inhibition from both adapter proteins [34]. Increases in glucose-stimulated insulin receptor signalling were more prominent in the skeletal muscle and white adipose tissue (WAT) of *Grb10* KO and *Grb10*:*Grb14* DKO mice and in liver of *Grb14* KO and *Grb10*:*Grb14* DKO mice. This suggests the additive effect is due to the two adapter proteins having a greater role in insulin receptor inhibition in different tissues and is consistent with the fact *Grb10* is lowly expressed in normal adult liver.

The physiological role of Grb7 is less clear, with no previous studies having described *Grb7* KO mice. In *in vitro* studies, Grb7 has mainly been associated with focal adhesion kinase-(FAK-) and ephrin B1-(ephB1-) mediated regulation of cell migration [35]. This is potentially an ancestral function since the RA and PH domains of the Grb7-family proteins are conserved in the *C. elegans* Mig-10 protein, which is required for embryonic neuronal migration. In addition, overexpression of human *GRB7* has been linked with progression of many cancer types, with evidence indicating it can affect diverse processes, including cell survival, proliferation, migration and invasion, mainly studied in cancer cell lines [36].

*GRB14* has been linked with several cancer types, with evidence that it promotes tumour progression in thyroid carcinoma [37] and glioblastoma [38], and may act as a tumour suppressor in hepatocellular carcinoma [39]. Similarly, there is evidence for *GRB10* having an oncogenic role in prostate carcinoma [40], glioma [41] and gastric cancer [42]. More in keeping with its growth inhibitory role during development there is convincing evidence *GRB10* acts as a tumour suppressor in at least two cases, that of clear cell renal cell carcinoma [43] and in tumours from a cancer-prone mouse model heterozygous for the Neurofibromatosis 1 (*Nf1*) gene [44].

Here we show that expression patterns of Grb7 and Grb14 each overlap with that of Grb10 during fetal development, and to a limited extent with each other. We include the first description of *Grb7* KO mice and address the potential for *Grb7* and *Grb14* to act redundantly with *Grb10* in the regulation of mouse fetal growth, including the expansion of liver due to excess lipid accumulation. Mean birth weights of *Grb7* KO and *Grb14* KO pups were similar to those of wild type littermates, whereas *Grb10* KO pups were significantly heavier, by approximately 30%, consistent with previous studies [17, 18, 23, 45]. Despite the overlapping expression patterns during development, tests for genetic interactions between *Grb10* and either *Grb7* or *Grb14* mice revealed no evidence of additive effects on fetal growth. In particular, there was no evidence from birth weights that either *Grb7*:*Grb10* DKO or *Grb10*:*Grb14* DKO pups were larger than *Grb10* single KO pups. Likewise, there was no evidence that either *Grb7* or *Grb14* contributed to the growth of most individual organs, with the exception that *Grb7* appeared to have a positive influence on fetal kidney growth. Initial characterization of *Grb7* KO adult mice revealed subtle changes in adipose deposition and glucose sensitivity, particularly in females, suggesting a role for *Grb7* as a physiological regulator of insulin signaling, which merits further study.

## Materials and Methods

### Mice

Derivation of KO mouse strains has already been described for *Grb10* (previously designated *Grb10Δ2-4*, full designation *Grb10^Gt(β-geo)1Ward^*) [17] and *Grb14* [33]. *Grb7* KO mice were transferred to the University of Bath facility as frozen embryos for this study after being generated at the University of Michigan, as shown in Figure 5A. In brief, homology arms from the 5’ portion of the gene (between a *Not*I restriction site in the upstream non-coding region and an *Nco*I site overlapping the translational start site in Exon 2) and from the 3’region (a *Bam*HI fragment in the non-coding sequence downstream of the final exon, Exon 15) were cloned into the pPNT vector [46] that includes a *neomycin* (*neo*) resistance gene cassette for positive selection and a herpes simplex virus (hsv) thymidine kinase (tk) gene for negative selection during targeting of embryonic stem (ES) cells. In targeted alleles the *neo* cassette replaced all the *Grb7* coding exons (Exons 2 to 15) to form a predictive null allele. Successful targeting was determined by Southern blotting, using standard techniques (as in [47]) of *Hind*III digested DNA from ES cells or mice using probes from within the 5’ homology arm (Probe A, PCR amplified using primers spanning Exon 1; 5’-TAGCACCTGCTGCTCAGT-3’ and 5’-GCAGCCTGAGAGGCTCCC-3’) and from immediately downstream of the 3’ homology arm (Probe B). For the wild type allele, Probe A detected a *Hind*III fragment of approximately 8.7kb and Probe B a 10.8kb fragment, whereas both probes detected a 15.3kb fragment for the KO allele. To generate experimental animals, first *Grb10^+/p^* males were crossed with either *Grb7^+/-^* or *Grb14^+/-^* females to produce *Grb7^+/-^*: *Grb10^+/p^* and *Grb10^+/p^*: *Grb14^+/-^* heterozygotes. Heterozygous *Grb7^+/-^*: *Grb10^+/p^* females were then crossed with *Grb7^+/-^*: *Grb10^+/+^* males to produce experimental offspring of six genotypes (Table 1A). Similarly, heterozygous *Grb10^+/p^*: *Grb14^+/-^* females were then crossed with *Grb14^+/-^*: *Grb10^+/+^* males to produce experimental offspring of six genotypes (Table 1B). In each case, offspring were sorted into four groups for analysis, as shown (Table 1). From these crosses embryos and placentae were collected on embryonic day e17.5, where e0.5 was the day on which a copulation plug was observed, or on the day of birth, designated post-natal day 1 (PN1). In addition, wild type (*Grb7^+/+^*) and *Grb7* KO (*Grb7^-/-^*) mice were compared at adult stages, and these were derived from separate intercrosses between *Grb7^+/-^* heterozygotes. Wild type littermates are considered the control group and single animals the biological replicate, noting that multiple litters were generated in each cross, with the aim of having enough of the least common genotypes for robust statistical analysis. All animals were maintained on a mixed inbred (C57BL/6J:CBA/CA) strain background and housed in individually ventilated cages under conditions of 13 hours light:11 hours darkness, including 30-minute periods of dim lighting to provide false dawn and dusk, a temperature of 21±2°C and relative humidity of 55±10%. Standard chow (CRM formula; Special Diets Services, Witham, Essex, UK) and water was available *ad libitum*. Experiments involving mice were subject to local ethical review by the University of Bath Animal Welfare and Ethics Review Board and carried out under licence from the United Kingdom Home Office. The manuscript has been written in accordance with the Animal Research: Reporting of In Vivo Experiments (ARRIVE) guidelines (https://arriveguidelines.org/).

**Table 1.** Genetic crosses used in the study, showing parent and offspring genotypes with their expected ratios. Crosses between. (A) *Grb7^+/−^:Grb10^+/p^*double heterozygous females and *Grb7^+/−^* heterozygous males, producing offspring of six genotypes, and (B) *Grb10^+/p^:Grb14^+/−^* double heterozygous females and *Grb10^+/p^:Grb14^+/−^* double heterozygous males, producing offspring of twelve genotypes, each in the indicated, expected Mendelian ratios. For ANOVA statistical analysis these six genotypes were used to form four groups, as indicated, since no difference was anticipated between animals differing only in their *Grb7^+/−^* and *Grb7^+/+^* or *Grb14^+/−^* and *Grb14^+/+^* allelic status. Similarly, due to the imprinted expression of *Grb10*, offspring inheriting a mutant copy of the paternal *Grb10* allele (*Grb10^+/p^*), having no growth phenotype, were not expected to differ from *Grb10* wild type (*Grb10^+/+^*), offspring, while double knockout (DKO) offspring inheriting mutations of both parental alleles (*Grb10^m/p^*) were expected to be indistinguishable from those inheriting a mutant copy of the normally active maternal *Grb10* allele (*Grb10^m/+^*).

| Parents | *Grb7*<sup>+/-</sup>:*Grb10*<sup>+p</sup> female x *Grb7*<sup>+/-</sup>:*Grb10*<sup>+/+</sup> male |  |  |  |  |  |
| --- | --- | --- | --- | --- | --- | --- |
| Offspring | *Grb7*<sup>+/+</sup>:<br>*Grb10*<sup>+/+</sup> | *Grb7*<sup>+/-</sup>:<br>*Grb10*<sup>+/+</sup> | *Grb7*<sup>-/-</sup>:<br>*Grb10*<sup>+/+</sup> | *Grb7*<sup>+/+</sup>:<br>*Grb10*<sup>m/+</sup> | *Grb7*<sup>+/-</sup>:<br>*Grb10*<sup>m/+</sup> | *Grb7*<sup>-/-</sup>:<br>*Grb10*<sup>m/+</sup> |
| Ratio | 1 | 2 | 1 | 1 | 2 | 1 |
| Group | Wild type |  | *Grb7* KO | *Grb10* KO |  | *Grb7*:*Grb10*<br>DKO |

B)
| Parents | *Grb10*<sup>+p</sup>: *Grb14*<sup>+/-</sup> female x *Grb10*<sup>+p</sup>: *Grb14*<sup>+/-</sup> male |  |  |  |  |  |
| --- | --- | --- | --- | --- | --- | --- |
| Offspring | *Grb10*<sup>+/+</sup>:<br>*Grb14*<sup>+/+</sup> | *Grb10*<sup>+/+</sup>:<br>*Grb14*<sup>+/-</sup> | *Grb10*<sup>+/+</sup>:<br>*Grb14*<sup>-/-</sup> | *Grb10*<sup>m/+</sup>:<br>*Grb14*<sup>+/+</sup> | *Grb10*<sup>m/+</sup>:<br>*Grb14*<sup>+/-</sup> | *Grb10*<sup>m/+</sup>:<br>*Grb14*<sup>-/-</sup> |
| Ratio | 1 | 2 | 1 | 1 | 2 | 1 |
|  | *Grb10*<sup>+p</sup>:<br>*Grb14*<sup>+/+</sup> | *Grb10*<sup>+p</sup>:<br>*Grb14*<sup>+/-</sup> | *Grb10*<sup>+p</sup>:<br>*Grb14*<sup>-/-</sup> | *Grb10*<sup>m/p</sup>:<br>*Grb14*<sup>+/+</sup> | *Grb10*<sup>m/p</sup>:<br>*Grb14*<sup>+/-</sup> | *Grb10*<sup>m/p</sup>:<br>*Grb14*<sup>-/-</sup> |
| Ratio | 1 | 2 | 1 | 1 | 2 | 1 |
| Group | Wild type |  | *Grb14* KO | *Grb10* KO |  | *Grb10*:<br>*Grb14*<br>DKO |

### PCR genotyping

For *Grb7* alleles, mice were genotyped by PCR using primers within the *neo* cassette (neoF, 5’-GCCCGGCATTCTGCACGCTT-3’; neoR, 5’-AGAGCAGCCGATTGTCTGTTGT-3’) to identify the targeted allele and from *Grb7* Exon 3 (deleted in the targeted allele, e3F, 5’-GTTTCAGGCAACCTCTCTGC-3’; e3R, 5’-TGGAGTCTCGAGGAAGCAAC-3’) to identify the wild type allele. Primers for genotyping *Grb14* alleles [33] were previously described, as were primers for *Grb10* alleles and PCR conditions [17].

### Tissue collection, histology and immunohistochemistry

Whole bodies (e17.5 and PN1) and PN1 organs were collected, any surface fluid removed from embryos or dissected organs by gently touching them onto absorbent paper, and weights obtained using a fine balance accurate to 4 decimal places (Sartorius BP61S). Paired organs (lungs and kidneys) were weighed together. Organs for histology were fixed by immersion in 4% (w/v) paraformaldehyde (PFA) in PBS at 4°C for 16-24 hours, then processed by machine (Leica TP1020) for wax embedding. Sections were cut at approximately 8-10 μm using a microtome (Leica Histocore Biocut), prior to staining with haematoxylin and eosin (H & E) using standard procedures [48]. Images were collected using a digital colour camera (Olympus SC50) and software (Olympus cellSens Entry), attached to a compound microscope (Nikon Eclipse E800). For immunohistochemistry, protocols were essentially as previously described [17], using primary antibodies specific for either Grb7 (rabbit polyclonal; Santa Cruz Biotechnology, CA, USA), at 1:200 dilution, or Grb14 (goat polyclonal; Santa Cruz Biotechnology, CA, USA) at 1:100 dilution. Biotin-conjugated anti-goat or anti-rabbit secondary antibodies were each applied at 1:500 dilution and incubated with Vector elite Reagent (Vector Laboratories, CA, USA) for 45 minutes, prior to developing with DAB reagent.

### Western blotting

Protein detection by western blot was carried out essentially as described [17] using 10μg of tissue lysate per sample, loaded on the basis of a bicinchoninic acid (BCA) assay (Pierce). Proteins were separated on 10% polyacrylamide gels and electroblotted to PVDF membranes. Membranes were probed with rabbit polyclonal primary antibodies specific for either Grb7 (Santa Cruz Biotechnology, CA, USA), used at 1:500 dilution, or for α-tubulin (Santa Cruz Biotechnology, CA, USA) used at 1:30,000 dilution, as a loading control. In each case, a peroxidase-conjugated goat anti-rabbit secondary antibody (Vector Laboratories, CA, USA) was used at 1:10,000 and proteins were visualised by enhanced chemiluminescence using the ECL-Plus system (GE Healthcare).

### Body composition, glucose measurements and food intake

Dual energy X-ray absorptiometry (DXA), using a PIXImus instrument (Lunar, Madison, WI, USA) with small-animal software [26] was used to collect data on adult mice for whole body, lean and fat mass, bone mineral content (BMC) and bone mineral density (BMD). The whole body and individual organs dissected from the same animals subject to DXA were also weighed, the organs on scales accurate to 4 decimal places (Sartorius BP61S). Glucose levels were obtained using a One-Touch ULTRA (Lifescan, CA) glucometer immediately following collection of whole blood during dissection of PN1 pups or from the tail vein of adult mice. Glucose tolerance tests and measurements of food intake were performed as previously described [26].

### Statistics

Chi-square tests were applied to determine whether the genotypes of experimental groups were present in the expected Mendelian ratios. Numerical data were usually subject to one-way analysis of variance (ANOVA), using a Kruskall-Wallis test with post-hoc Dunn’s test to determine p-values between groups. This test allowed us to detect significant differences associated with either of the single knockout groups in each set of progeny as well as any significant interaction between them. This relatively conservative non-parametric test was chosen because in some experiments one or more genotype group was represented by a small samples size (n=<5). Where only two groups were being compared a student’s t-test was applied, except in instances where the variance of the two groups were significantly different, in which case a non-parametric Mann Whitney test was used. All statistical tests were applied using GraphPad Prism (v10 GraphPad, La Jolla, CA, USA) software. Graphs show arithmetic means ± standard error of the mean (SEM). Differences with p-values of <0.05 were considered statistically significant.

## Results

### Expression of Grb7 and Grb14 in the embryo and in adult pancreas

#### Tissue distribution of Grb7 in the e14.5 mouse embryo

To date, no extensive developmental expression pattern of mouse Grb7 has been reported, with available information based on Northern blots of RNA prepared from homogenised adult tissues [3, 49]. One exception is a study that included mRNA in situ hybridisation data, providing spatial information only for fetal mouse lung, gut and kidney [50]. Consequently, we sought to characterise the expression pattern of Grb7 during embryonic mouse development by immunohistochemistry using an antibody raised against the N-terminus of Grb7. Analysis of histological sections from e14.5 embryos revealed a pattern of Grb7 expression in tissues of mesodermal and endodermal origin. Grb7 was readily detected in developing liver, pituitary, inner ear, nasal epithelium and tooth primordia, along with strong staining of epithelial structures, including the epidermis (Figure 1A), within the submandibular salivary gland (Figure 1B), kidney tubules (Figure 1C), bronchi within the lung (Figure 1D), lining of the gut (Figure 1E) and stomach (Figure 1A). There was also intense staining in the endocrine pancreas (Figure 1F), with lower levels of expression evident in the pericardium and ossified cartilage, for example in ribs (Figure 1D), as well as in the adrenal cortex (Figure 1C). Discrete staining was observed in the endocrine component of the developing pancreas (Figure 1E). Notably, Grb7 was absent from the CNS and different muscle types, including skeletal muscle, smooth muscle of the gut, diaphragm and cardiac muscle.

**Figure 1.**
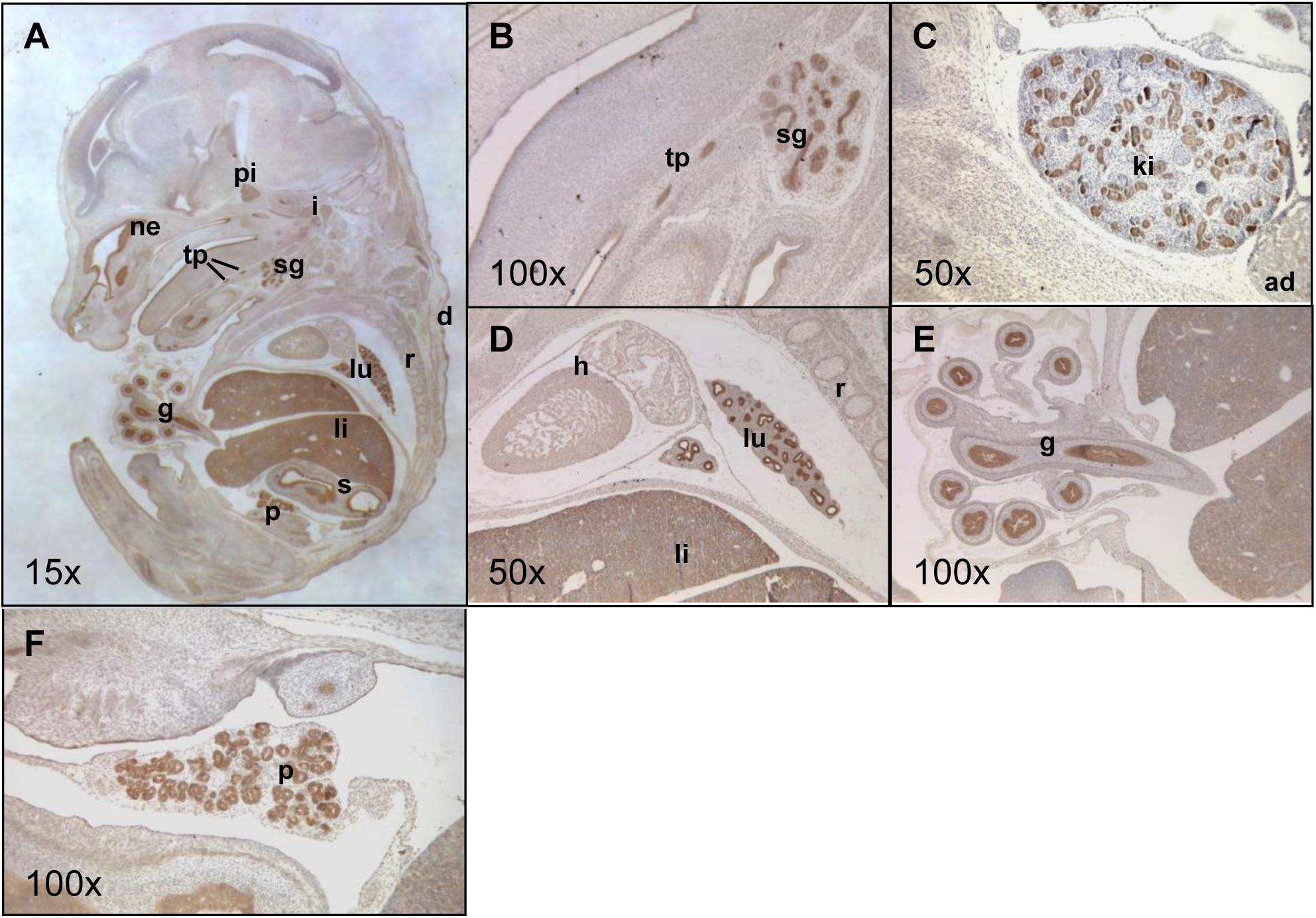
Grb7 expression in the mouse embryo visualised by immunohistochemistry on wild type e14.5 paraffin sections. Representative mid-sagittal sections were chosen to display a wide range of tissues. (A) Whole embryo with expression highlighted in dermis (d), gut (g), inner ear (i), liver (li), lung (lu), nasal epithelium (ne), pancreas (p), pituitary (pi), ribs (r, ossified cartilage), stomach (s), salivary gland (sg) and tooth primordia (tp); (B) Expression in submandibular gland and tooth primordia; (C) Expression in kidney (ki) and adrenal gland (ad); (D) Expression in lung and liver, but not heart (h); (E) Expression in epithelial lining of mid– and hind-gut; (F) Endocrine pancreas. Brown staining is indicative of Grb7 expression, magnifications as indicated.

#### Tissue distribution of Grb14 in the e14.5 mouse embryo

Previous studies of Grb14 expression in the mouse have been restricted to methods for bulk tissue analysis such as RT-PCR, Northern and Western blotting (e.g. [33]). Consequently, we again employed immunohistochemistry to characterise the tissue distribution of Grb14 in e14.5 embryo sections, where possible using sections adjacent to those used to probe for Grb7, to facilitate direct comparison. An antibody raised against the N-terminus of Grb14 was used and revealed a distinct pattern of expression (Figure 2A). In the brain, Grb14 was restricted to the apical surface of the choroid plexus epithelial layer (Figure 2B). Grb14 protein was detected at high levels in the ossifying cartilage, including, vertebrae (Figure 2C) and ribs (Figure 2D), as well as in various in skeletal muscles, including intercostals (Figure 2D), diaphragm and tongue (Figure 2A). High levels were additionally found in the tooth primordia, throughout the cardiac muscle and dermis. At lower levels, Grb14 was found in the developing bronchi of the lungs (Figure 2D), throughout the pancreatic primordium (Figure 2E), and the epithelial lining of the midgut and stomach, (Figure 2A). Grb14 was not detected in the pituitary, pericardium, smooth muscle of the gut, kidney or adrenal gland. A summary of the expression findings for Grb7 and Grb14 at e14.5, in comparison with the previously described pattern for Grb10 [17, 18, 45] is provided (Table 2). Having observed expression of all three adaptor proteins in the developing pancreas, we also examined their expression in the adult organ. Grb7, Grb10 and Grb14 proteins were each readily detected throughout the endocrine cells of the islets of Langerhans (Figure 3).

**Figure 2.**
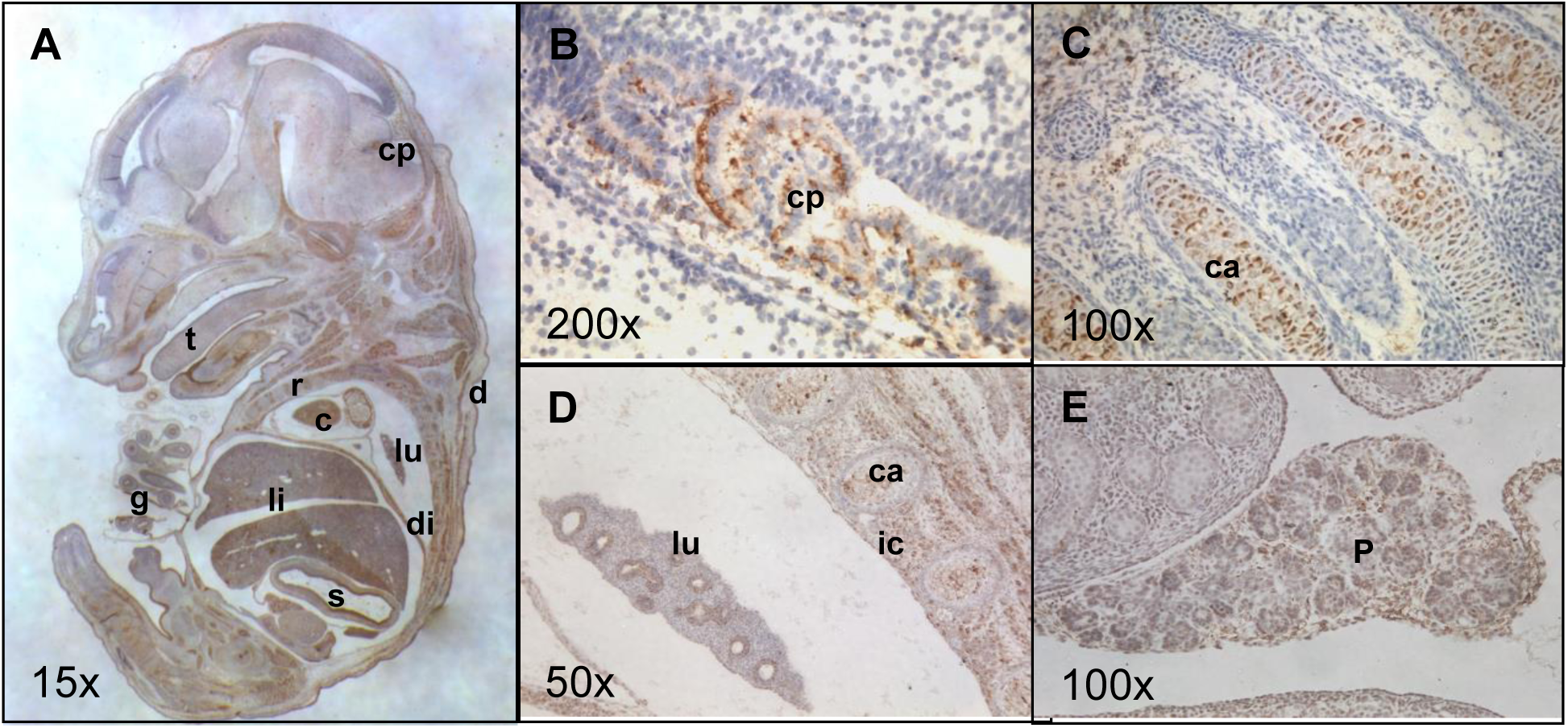
Grb14 expression in the mouse embryo visualised by immunohistochemistry on wild type e14.5 paraffin sections. Representative mid-sagittal sections were chosen to display a wide range of tissues. (A) Whole embryo with expression highlighted in cardiac muscle (c), choroid plexus (cp), dermis (d), diaphragm (di), gut (g), lungs (lu), liver (li), ribs (r, ossifying cartilage), stomach (s) and tooth primordia (tp); (B) Expression in choroid plexus; (C) Expression in ossifying cartilage (ca) of vertebrae; (D) Expression in lung, ossifying cartilage of ribs and intercostal muscle (ic); (E) Expression in pancreas (p). Brown staining is indicative of Grb14 expression, magnifications as indicated.

**Figure 3.**
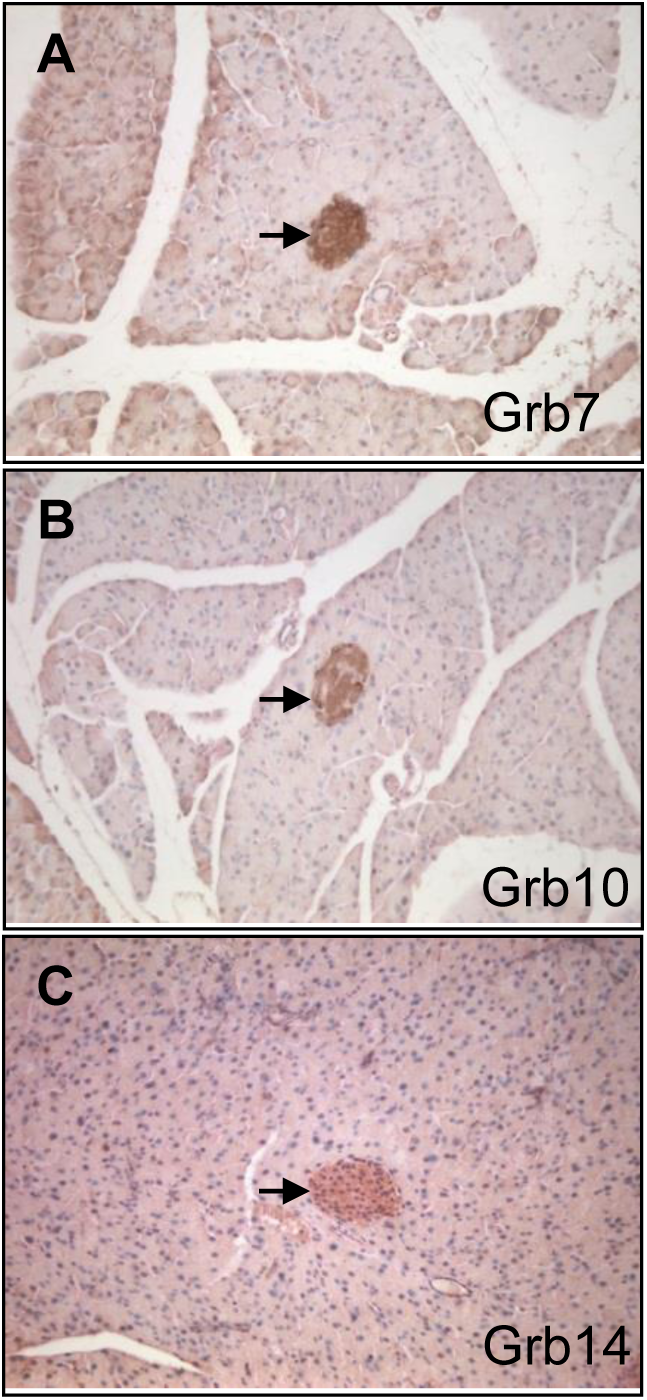
Expression of Grb7 family members in the PN21 mouse pancreas visualised by immunohistochemistry on paraffin sections with antibodies specific for (A) Grb7; (B) Grb10; and C) Grb14. Brown staining is indicative of protein expression. Arrows indicate endocrine area (pancreatic islets); magnification 100x.

**Table 2.** Grb7, Grb10 and Grb14 distribution in the developing mouse embryo. Expression determined by immunohistochemistry on wild type embryo sections at e14.5. Expression is compared in the listed tissues with notes on the specific localisation of each protein. An indication of relative expression level (E) is provided, in which levels have been judged separately for each antibody through comparison of tissues in the same sections (H = high, M = medium, L = low, V = very low). Grb10 is expressed in the CNS exclusively from the paternally inherited allele (indicated *) and in all other sites exclusively or primarily from the maternally inherited allele. Ch plexus = choroid plexus; gl = gland; Nasal ep = nasal epithelium; Sk muscle = skeletal muscle; Submandib = submandibular salivary gland; Tooth prim = tooth primordium.

| Tissue | Grb7 |  | Grb10 |  | Grb14 |  |
| --- | --- | --- | --- | --- | --- | --- |
|  | Details | E | Details | E | Details | E |
| CNS | No | - | Yes*, ventral spinal cord, mid-brain | M | No | - |
| Ch plexus | inconclusive | - | Yes, epithelium | H | Yes, apical epithelial surface | H |
| Pituitary | Yes | H | Yes | M | No | - |
| Submandib | Yes, epithelium | H | No | - | No | - |
| Inner ear | Yes, epithelium | H | Yes | M | Yes | M |
| Nasal ep | Yes | H | Yes | M | Yes | M |
| Tooth prim | Yes | M | Yes | M | Yes | M |
| Cartilage | Yes, ossified | V | Yes | H | Yes, ossifying | H |
| Sk muscle | No | - | Yes, widely | H | Yes, intercostal,<br>diaphragm | H |
| Heart | Yes, Pericardium | L | Yes, Cardiac<br>muscle | M | Yes, Cardiac<br>muscle | H |
| Lung | Yes, epithelial | H | Yes, epithelial | H | Yes, epithelial | M |
| liver | Yes | H | Yes | H | Yes | M |
| Pancreas | Yes, endocrine | H | Yes, endocrine | H | Yes, endocrine | M |
| Kidney | Yes, epithelial | H | Yes, epithelial | H | No | - |
| Adrenal gl | Yes, cortex | L | Yes, cortex | M | No | - |
| Stomach | Yes, epithelial | H | Yes, epithelial | H | Yes, epithelial | L |
| Lower gut | Yes, epithelial | H | Yes, epithelial | H | Yes, epithelial | V |
| Epidermis | No | - | Yes | M | Yes | M |

### Functional overlap between Grb10 and Grb14 in the regulation of fetal growth

#### *Grb10* KO x *Grb14* KO offspring PN1 body mass and blood glucose levels

Progeny of crosses between *Grb10^+/p^*:*Grb14^+/−^*females and *Grb10^+/−^*:*Grb14^+/−^* males were collected at PN1 for body and organ weight analysis (Figure 4). A Chi-squared test indicated that offspring of the twelve anticipated genotypes were present at the expected frequencies (p=0.8629) (Supp. Table 1A). Progeny of the twelve genotypes were reduced to four groups by treating the following pairs of offspring as equivalent; *Grb10^+/+^* and *Grb10^+/p^*; *Grb10^m/+^* and *Grb10^m/p^*; and *Grb14^+/−^* and *Grb14^+/+^* (Table 1B). Pooling allowed us to strengthen statistical analyses, while simplifying data analysis and presentation, without materially affecting the analysis. Mean body and organ weights are summarised in Table 3A. Compared to wild type controls (mean weight 1.3827±0.0275g), *Grb10* KO pups (1.8306g±0.0430g) were approximately 32% larger (p<0.0001), a size difference consistent with that seen previously [17, 18, 22, 23] and in keeping with the role for *Grb10* as a fetal growth inhibitor (Figure 4A). In contrast, *Grb14* KO pups (1.3657±0.0288g) were not significantly different to wild type with a mean weight only 1% smaller. *Grb10*:*Grb14* DKO animals (1.6003±0.0789g) were intermediate in size between wild type and *Grb10* KOs (16% larger than wild type), without being significantly different to either. This was unexpected since the most likely predicted outcomes were either that the DKO pups would be very similar to *Grb10* KO single mutants, indicating no redundancy in growth regulation, or DKOs would be even larger than *Grb10* KOs, indicating functional redundancy between the two adaptor proteins as fetal growth inhibitors. The observation that loss of *Grb14* should have a negative influence on growth only in the absence of *Grb10* is difficult to explain at a mechanistic level. To investigate whether Grb10 and Grb14 might influence circulating glucose levels in neonates through combined actions on the Insr, we also measured blood glucose at PN1 (Figure 4B). Mean values for wild type (3.0±0.2 mM), *Grb10* KO (3.3±0.8 mM), *Grb14* KO (3.5±0.5 mM) and *Grb10*:*Grb14* DKO (3.4±0.6 mM) were all very similar and no significant differences were found between any of the genotypes.

**Figure 4.**
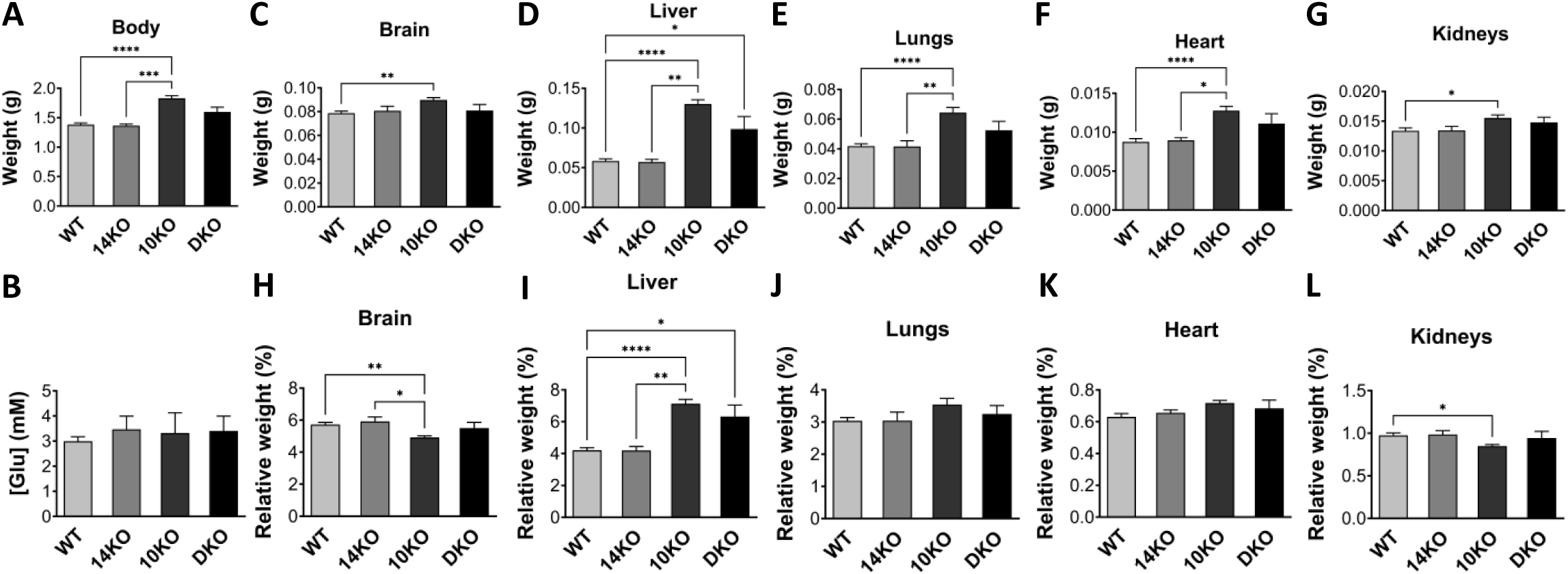
Weights of whole body and selected dissected organs, with blood glucose levels, at PN1 from progeny of crosses between *Grb10* KO and *Grb14* KO mice. Data were pooled into four groups for analysis as described in the Methods; wild type (WT), *Grb10* KO (10KO), *Grb14* KO (14KO) and *Grb10*:*Grb14* double knockout (DKO). For each of the four offspring genotype groups, data are shown for (A) Body weight; and (B) Blood glucose concentration ([Glu]). In addition, actual weights of (C) Brain; (D) Liver; (E) Lungs; (F) Heart; and (G) Kidneys; are shown above the relative weights of the same organs, expressed as a percentage of body mass (H-L). Values represent means and SEM, tested by one-way ANOVA using Kruskal-Wallis and Dunn’s post hoc statistical tests. Sample sizes were, for total body and all organs, WT n=25, *Grb10* KO n=13, *Grb14* KO n=7, *Grb10*:*Grb14* DKO n=7; glucose levels, WT n=18, *Grb10* KO n=9, *Grb14* KO n=6, *Grb10*:*Grb14* DKO n=4. Asterisks indicate *p*-values, *p <0.05, **p <0.01, ***p <0.001, ****p<0.0001.

**Table 3.**
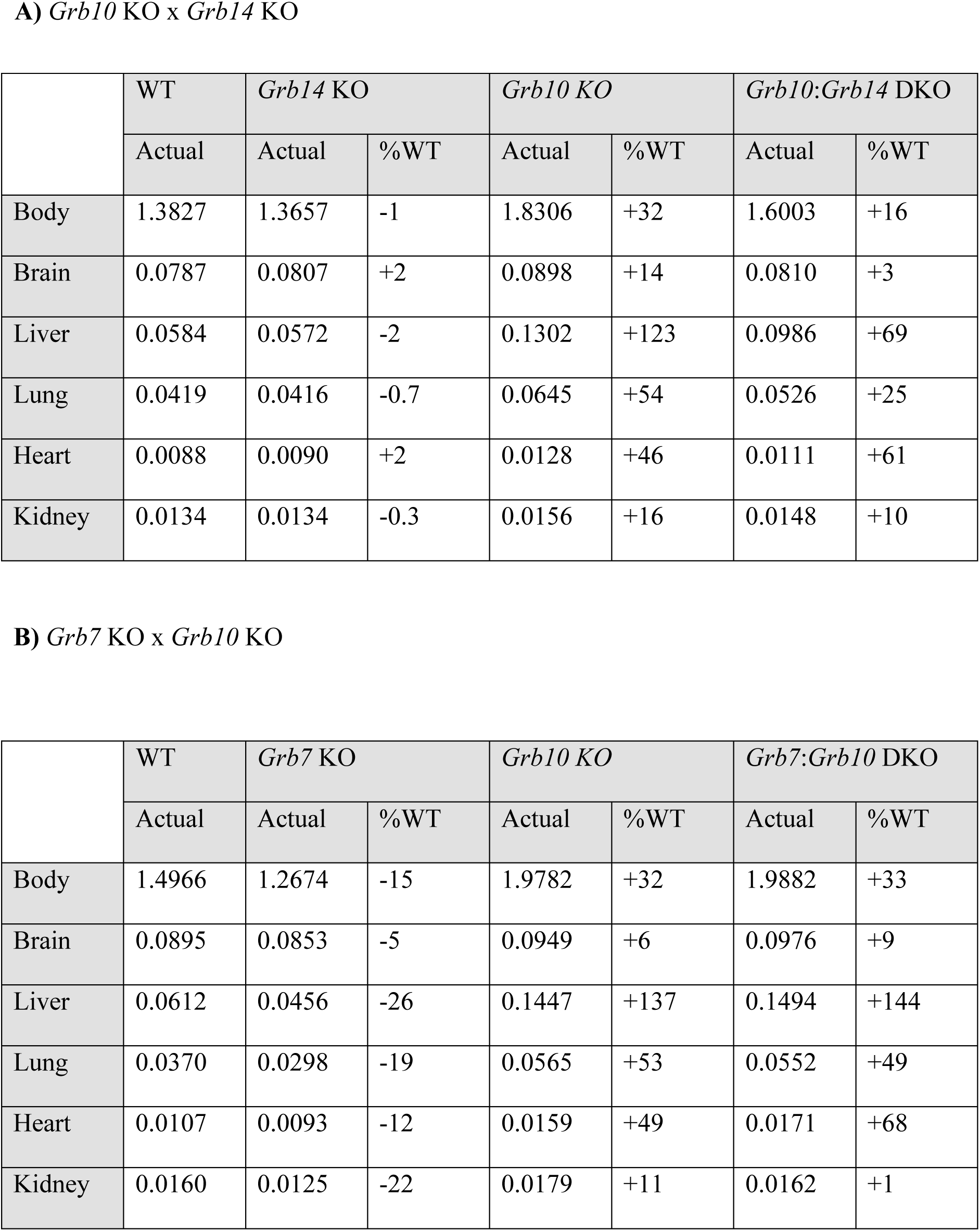
Summary of PN1 body and organ weight data for progeny of crosses between. (A) the *Grb10* KO and *Grb14* KO strains, and (B) the *Grb7* KO and *Grb10* KO strains. Mean weights are shown for each genotype together with changes relative to wild type (%WT) for each mutant genotype.

|  | WT | *Grb14* KO |  | *Grb10* KO |  | *Grb10:Grb14* DKO |  |
| --- | --- | --- | --- | --- | --- | --- | --- |
|  | Actual | Actual | %WT | Actual | %WT | Actual | %WT |
| Body | 1.3827 | 1.3657 | -1 | 1.8306 | +32 | 1.6003 | +16 |
| Brain | 0.0787 | 0.0807 | +2 | 0.0898 | +14 | 0.0810 | +3 |
| Liver | 0.0584 | 0.0572 | -2 | 0.1302 | +123 | 0.0986 | +69 |
| Lung | 0.0419 | 0.0416 | -0.7 | 0.0645 | +54 | 0.0526 | +25 |
| Heart | 0.0088 | 0.0090 | +2 | 0.0128 | +46 | 0.0111 | +61 |
| Kidney | 0.0134 | 0.0134 | -0.3 | 0.0156 | +16 | 0.0148 | +10 |

**B) *Grb7* KO x *Grb10* KO**
|  | WT | *Grb7* KO |  | *Grb10* KO |  | *Grb7:Grb10* DKO |  |
| --- | --- | --- | --- | --- | --- | --- | --- |
|  | Actual | Actual | %WT | Actual | %WT | Actual | %WT |
| Body | 1.4966 | 1.2674 | -15 | 1.9782 | +32 | 1.9882 | +33 |
| Brain | 0.0895 | 0.0853 | -5 | 0.0949 | +6 | 0.0976 | +9 |
| Liver | 0.0612 | 0.0456 | -26 | 0.1447 | +137 | 0.1494 | +144 |
| Lung | 0.0370 | 0.0298 | -19 | 0.0565 | +53 | 0.0552 | +49 |
| Heart | 0.0107 | 0.0093 | -12 | 0.0159 | +49 | 0.0171 | +68 |
| Kidney | 0.0160 | 0.0125 | -22 | 0.0179 | +11 | 0.0162 | +1 |

#### *Grb10* KO x *Grb14* KO offspring PN1 organ mass

To assess body proportions and potential selective effects on individual organs, brain, liver, lungs, heart and kidneys were dissected at PN1, with weights analysed directly (Figure 4C-G) and as a percentage of total body weight (Figure 4H-L). First, as seen in our previous studies the brain from *Grb10* KO pups was spared from the more dramatic overgrowth of the body, being only 14% (p<0.01) larger than wild type (0.08987±0.0020 versus 0.0787±0.0018g) (Figure 4C). In contrast, brains from *Grb14* KO (0.0807±0.0039g) and *Grb10*:*Grb14* DKO (0.0810±0.0050g) pups did not significantly differ from each other or from brains of the other two genotypes. This meant that while *Grb14* KO and *Grb10*:*Grb14* DKO brains did not differ significantly from wild type proportions, the *Grb10* KO brain was small (p<0.01) relative to an enlarged body (Figure 4H). A similar sparing effect was seen in *Grb10* KO kidneys (0.0156±0.0005g), which were only 16% larger (p<0.05) than wild type (0.0134±0.0047g) (Figure 4G) and disproportionately small (p<0.05) within the 32% bigger body (Figure 4L). Kidneys from *Grb14* KO (0.0134±0.0007g) pups were near identical to wild type and those from *Grb10*:*Grb14* DKO (0.0148±0.0009g) 10% larger, though not significantly different.

In contrast to brain and kidney, *Grb10* KO liver (0.1302±0.0054g) was 123% larger (p<0.0001) than the wild type liver (0.0584±0.0028g)(Figure 4D). *Grb14* KO liver (0.0572±0.0034g) was indistinguishable from wild type, while *Grb10*:*Grb14* DKO liver (0.0986±0.0036g) was 69% larger than wild type (p<0.05) and was not significantly different to *Grb10* KO liver. This meant the livers of both *Grb10* KO (p<0.0001) and *Grb10*:*Grb14* DKO (p<0.05) were disproportionately enlarged (Figure 4I) was not. Similarly, compared to wild type (0.0419±0.0015g), *Grb10* KO lungs (0.0645±0.0034g) were 54% larger (p<0.0001) and *Grb10*:*Grb14* DKO (0.0526±0.0059g) 25% larger, whereas *Grb14* KO lungs (0.0416±0.0004g) were almost identical to wild type (Figure 4E). In all cases the single KO and DKO mutant remained proportionate with body weight (Figure 4J). In the case of heart, only *Grb10* KO (0.1278±0.00056g) differed significantly (p<0.0001) from wild type (0.0088±0.0004g), being 46% larger, whereas *Grb14* KO heart (0.0090±0.0003g) were very similar, at only 2% smaller, and *Grb10*:*Grb14* DKO (0.0111±0.0013g) 27% larger (Figure 4F). In all cases the heart was proportionate with total body weight (Figure 4K).

#### Generation of a Grb7 knockout mouse and loss of protein expression

A *Grb7* KO mouse was generated using the strategy outlined in the Methods and Figure 5A. Successfully targeted ES cell clones were identified by Southern blotting of *Hind*III digested DNA, using ^32^P-labelled probes to detect bands of distinct sizes for the wild type and targeted alleles from the 5’– and 3’-ends of the gene (Figure 5B). Three successfully targeted ES cell clones were identified, one of which went on to produce a chimera that faithfully transmitted the genetic modification to subsequent generations. Since all of the coding exons were deleted from the targeted allele, all protein expression should be lost in *Grb7^-/-^* animals. This was tested by Western blotting protein extracts from liver and kidney of 12 week old animals. On blots probed with an anti-Grb7 antibody a single species of approximately 65 kDa, consistent with the predicted size of Grb7, was readily detected in tissues from *Grb7^+/+^* animals and was absent in *Grb7^-/-^* animals (Figure 5C), even after prolonged exposure (not shown). An α-tubulin specific antibody used as a loading control recognised a single 50 kDa band, readily detected in all samples.

**Figure 5.**
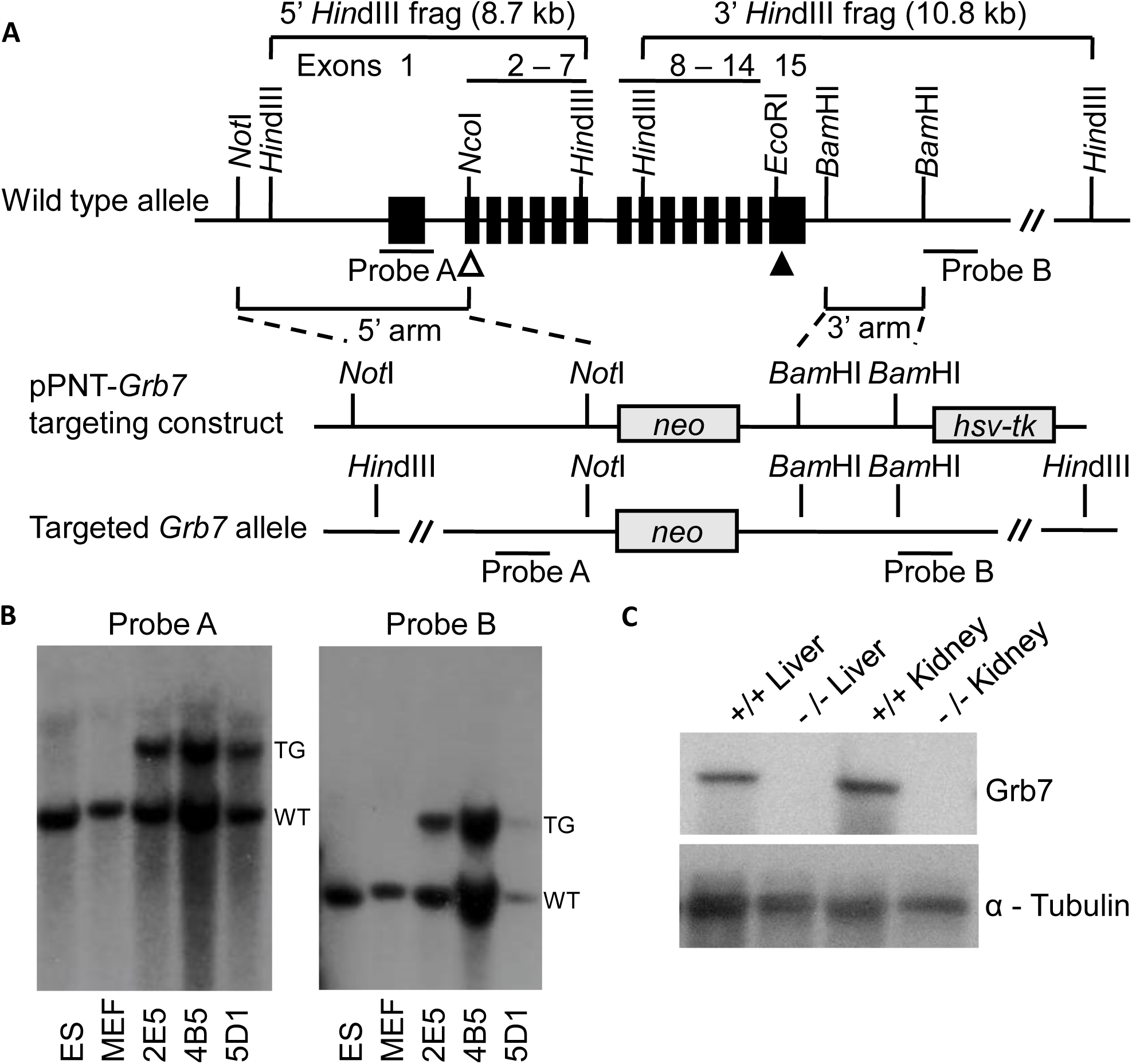
Generation of a *Grb7* KO mouse strain designed as a null mutation, lacking all the protein coding sequence. (A) The wild type *Grb7* locus (top), showing exons (filled boxes numbered 1-15), translational start (open triangle) and stop (filled triangle) codons, plus selected restriction enzyme sites. Note that the translational start (ATG) codon lies within a *Nco*I restriction site (CC<u>ATG</u>G) used as part of the cloning strategy. Regions used to direct homologous recombination in ES cells (5’ arm and 3’ arm) and also probes used to confirm correct targeting are indicated to either side of the coding exons. The homologous arms were cloned into the pPNT vector to form the pPNT-*Grb7* targeting construct (middle), in which the coding exons have been replaced with a *neomycin* (*neo*) resistance gene cassette for positive selection of ES cells stably incorporating the construct. The targeting construct also includes a herpes simplex virus thymidine kinase (*hsv-tk*) gene, located outside the homologous regions that is typically retained at sites of random integration, and was used to enrich for targeting events by negative selection. Consequently, the correctly targeted allele (bottom) retains the *neo* but not the *hsv-tk* gene. (B) Southern blot analysis of *Hind*III digested DNA from wild type ES cells (ES) or primary mouse embryonic fibroblasts (MEF) alongside three successfully targeted ES cell clones (2E5, 4B5 and 5D1). In the targeted (TG) allele, loss of the sequence between the homologous arms alters the distance between restriction enzyme sites, compared with the wild type (WT) allele. Consequently, in the wild type allele Probe A recognises a 5’ 8.7kb fragment and Probe B a 3’ 10.8kb fragment, whereas both probes detect a15.3kb fragment for the targeted allele. (C) Western blot analysis of protein extracts from adult kidney and liver derived from *Grb7* wild type (+/+) and *Grb7* KO (-/-) homozygous mice following establishment of a true breeding line. The blot was probed with an antibody specific for Grb7, that readily detects a species of approximately the correct size (65kDa) in wild type but not *Grb7* KO samples, whereas an antibody specific for α-tubulin (50kDa) was detected equally in both.

### The effect of *Grb7* on fetal growth is restricted to a positive influence on kidney growth

#### *Grb7* KO x *Grb10* KO offspring PN1 body mass and blood glucose levels

Progeny of crosses between *Grb7^+/−^*:*Grb10^+/p^* females and *Grb7^+/−^*:*Grb10^+/+^* males were collected at PN1 for body and organ weight analysis (Figure 6). A Chi-squared test indicated that offspring of the six anticipated genotypes were present at the expected frequencies (Supp. Table 2A) (p=0.3653). Progeny with six genotypes were reduced to four groups by pooling *Grb7^+/−^*:*Grb10^+/+^*with *Grb7^+/+^:Grb10^+/+^* (wild type group) and *Grb7^+/+^*:*Grb10^m/+^* with *Grb7^+/−^*:*Grb10^m/+^* (*Grb10* KO group), for comparison with the two remaining genotypes, *Grb7^-/-^:Grb10^+/+^* (*Grb7* KO group) and *Grb7^-/-^*:*Grb10^m/+^*(*Grb7*:*Grb10* DKO group) (Table 1A). Mean body and organ weights are summarised in Table 3B. In comparison with wild type (1.4966±0.0305g), *Grb7* KO pups (1.2674g±0.0596) were 15% smaller, which was not significantly different. In contrast, *Grb10* KO (1.9782±0.0237g) and *Grb7*:*Grb10* DKO animals (1.9882±0.0535g) were each significantly larger than wild type, by 32% (p<0.0001) and 33% (p<0.0001) respectively, and near identical to each other (Figure 6A). This shws that loss of *Grb7* has little or no effect on prenatal growth, either alone or in combination with *Grb10*. To investigate whether Grb7 and Grb10 might influence circulating glucose levels in neonates through combined actions on the Insr, we measured blood glucose at PN1 (Figure 6B). The mean values for wild type (3.0mM), *Grb7* KO (3.4mM) and *Grb10* KO (2.9mM) glucose levels were not significantly different, whereas that for *Grb7*:*Grb10* DKO (2.2mM) was significantly lower than those for both wild type (p<0.05) and *Grb7* KO (p<0.01), lending support to the idea that Grb7 and Grb10 function redundantly to regulate Insr-regulated glucose levels.

**Figure 6.**
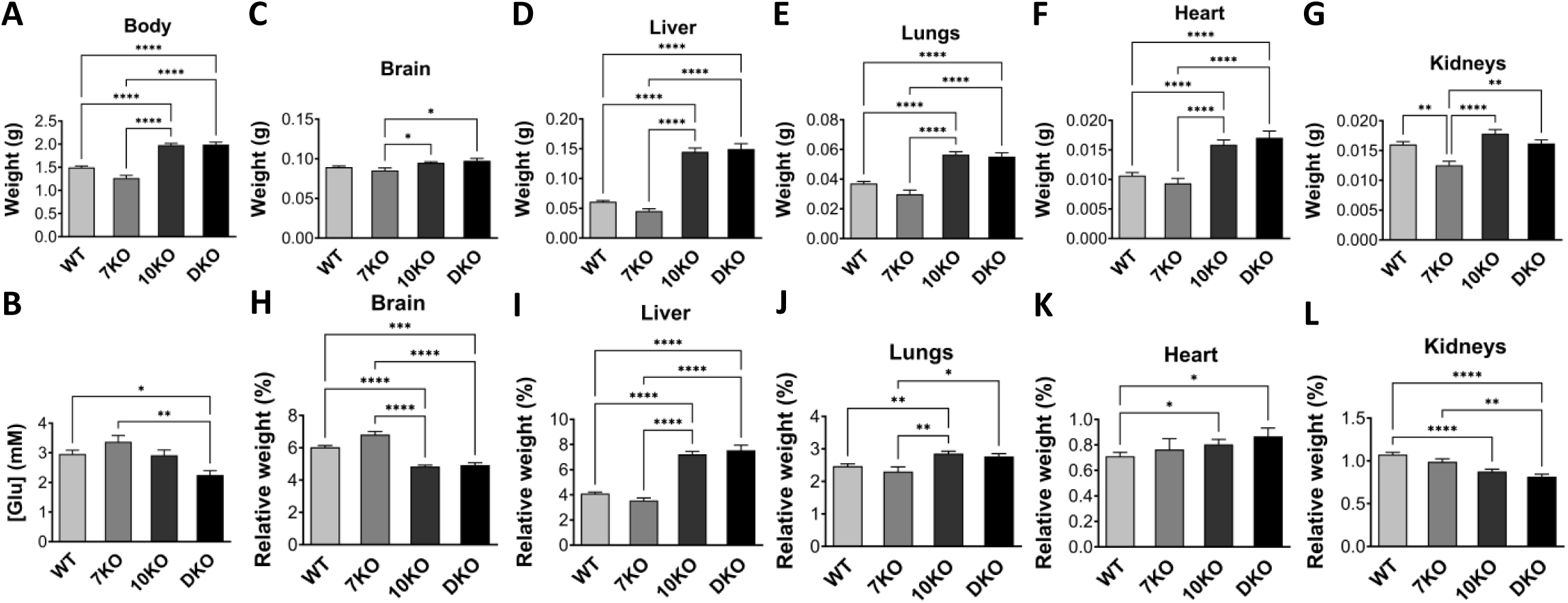
Weights of whole body and selected dissected organs, with blood glucose levels, at PN1 from progeny of crosses between *Grb7* KO and *Grb10* KO mice. Data were pooled into four groups for analysis as described in the Methods; wild type (WT), *Grb7* KO (7KO), *Grb10* KO (10KO) and *Grb7*:*Grb10* double knockout (DKO). For each of the four offspring genotype groups, data are shown for, (A) Body weight; and (B) Blood glucose concentration ([glu]). In addition, actual weights of (C) Brain; (D) Liver; (E) Lungs; F) Heart and G) Kidneys are shown above the relative weights of the same organs, expressed as a percentage of body mass (H-L). Values represent means and SEM, tested by one-way ANOVA using Kruskal-Wallis and Dunn’s post hoc statistical tests. Sample sizes were, for body and brain, WT n=42, *Grb7* KO n=14, *Grb10* KO n=49, *Grb7*:*Grb10* DKO n=15; kidneys and heart, WT n=42, *Grb7* KO n=14, *Grb10* KO n=48, *Grb7*:*Grb10* DKO n=15, liver, WT n=42, *Grb7* KO n=14, *Grb10* KO n=49, *Grb7*:*Grb10* DKO n=14; lungs, WT n=41, *Grb7* KO n=14, *Grb10* KO n=49, *Grb7*:*Grb10* DKO n=15; glucose levels, WT n=24, *Grb7* KO n=7, *Grb10* KO n=27, *Grb7*:*Grb10* DKO n=10. Asterisks indicate *p*– values, *p <0.05, **p <0.01, ****p<0.0001.

#### *Grb7* KO x *Grb10* KO offspring PN1 organ mass

Next, to assess the potential for Grb7 and Grb10 to selectively influence the growth of specific organs, either alone or in combination, the same selection of organs as before were dissected at PN1 and weights analysed directly (Figure 6C-G) and as a percentage of total body weight (Figure 6H-L). As in the previous cross, the *Grb10* KO brain (0.0949±0.0013g) was only slightly enlarged, in this case by 6% compared to wild type (0.0895±0.0015g) and was therefore spared from being overgrown to the extent of the whole body (Figure 6C). The brain of *Grb7*:*Grb10* DKO (0.0976±0.0029g) pups was similarly 9% larger, while the brain from *Grb7* KO (0.0853±0.0031g) pups was 5% smaller. Although none of the brains from mutant mice were significantly different to wild type, both *Grb10* KO and *Grb7*:*Grb10* DKO were significantly larger than *Grb7* KO brain (p<0.05 in each case). Consequently, *Grb10* KO and *Grb7*:*Grb10* DKO brains were each disproportionately small relative to both wild type (p<0.0001 and p<0.001, respectively) and *Grb7* KO (p<0.0001 in each case) brain (Figure 6H). A similar sparing effect was seen in kidney, for *Grb10* KO (0.0179±0.0068g), which were some 11% larger than wild type (0.0160±0.0005g). In striking contrast, *Grb7* KO kidneys (0.0125±0.0007g) were smaller by 22% (p<0.01), while *Grb7*:*Grb10* DKO (0.0162±0.0006g) were intermediate in size at just 1% larger than wild type (Figure 6G). Due to the small increases in size compared to wild type, kidneys from *Grb10* KO and *Grb7*:*Grb10* DKO were disproportionately small (p<0.0001 in each case) relative to wild type body proportions, whereas *Grb7* KO kidneys were proportionate (Figure 6L).

As before, *Grb10* KO liver (0.1447±0.0063g) was dramatically enlarged, this time by 137% (p<0.0001) compared with wild type (0.0612±0.0019g) (Figure 6D). *Grb7*:*Grb10* DKO liver (0.1494±0.0090g) was similarly 144% enlarged (p<0.0001), whereas *Grb7* KO liver (0.0456±0.0037g) was 26% smaller than wild type, though this difference was not statistically significant. This meant that both *Grb10* KO and *Grb7*:*Grb10* DKO livers were disproportionately enlarged compared to both wild type and *Grb7* KO (p<0.0001 in all cases), while *Grb7* KO livers did not deviate significantly from proportionality (Figure 6I).

The remaining organs followed a similar pattern to that of the body. Compared to wild type (0.0370±0.0014g), *Grb10* KO lungs (0.0565±0.0019g) were 53% larger (p<0.0001) and *Grb7*:*Grb10* DKO (0.0552±0.0025g) 49% larger (p<0.0001). *Grb7* KO (0.0298±0.0028g) lungs were 19% smaller, but not significantly different to wild type (Figure 6E). Relative to body weight, *Grb10* KO lungs were disproportionately enlarged compared to both wild type (P<0.01) and Grb7 KO (P<0.01). *Grb7*:*Grb10* DKO lungs were disproportionate only compared to Grb7 KO (p<0.05), while *Grb7* KO were proportionate (Figure 6J).

Similarly, both *Grb10* KO (0.0159±0.0008g) and *Grb7*:*Grb10* DKO hearts (0.0171±0.0011g) were significantly enlarged, being 49% (p<0.0001) and 60% (p<0.0001) bigger than wild type (0.0107±0.0005g), with *Grb7* KO (0.0093±0.0084g) closer to wild type at 12% smaller (Figure 6F). Both *Grb10* KO and *Grb7*:*Grb10* DKO hearts were marginally disproportionately enlarged (p<0.05 in each case) and *Grb7* KO proportionate (Figure 6K).

#### *Grb7* KO x *Grb10* KO offspring e17.5 embryo and placenta

To further investigate the potential for interaction between *Grb7* and *Grb10* to regulate growth, including by acting within the placenta, we analysed weights of the whole embryo and placenta at e17.5 (Figure 7). We chose a time-point late in gestation when any size differences between conceptuses of different genotypes would be relatively large and the placenta maximal in size [51, 52]. A Chi-squared test indicated that offspring of the six anticipated genotypes (Table 1A) were present at the expected frequencies (Supp. Table 2B) (p=0.7303) For the embryo, the pattern of weight differences was very similar to that of PN1 pups. *Grb10* KO embryos (1.2656±0.0387g) were 37% larger (p<0.0001) and *Grb7*:*Grb10* DKO (1.3117±0.0614g) 42% larger (p<0.05) than wild type (0.9624±0.0389g). *Grb7* KO (0.8724±0.0212g) embryos were 6% smaller but not significantly different to wild type (Figure 7A).

**Figure 7.**
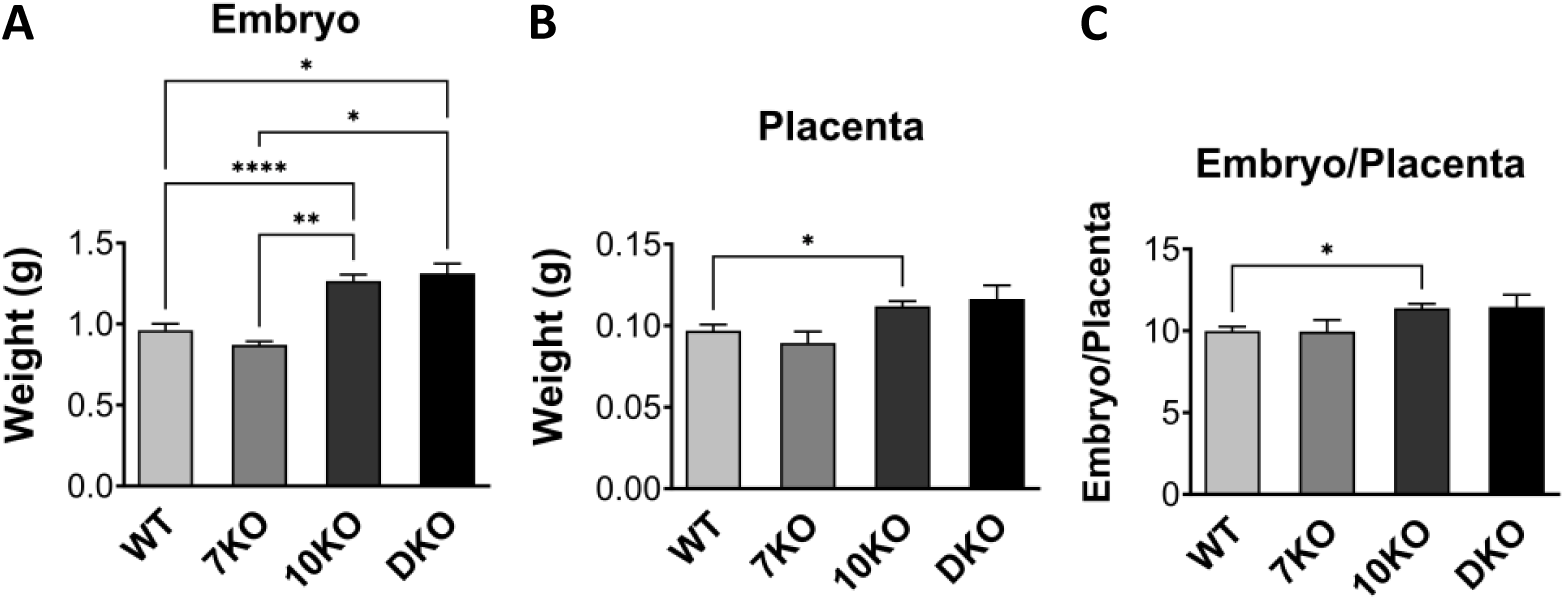
Weight analysis of e17.5 conceptuses from crosses between *Grb7* KO and *Grb10* KO mice. Data were pooled into four groups for analysis as described in the Methods; wild type (WT), *Grb7* KO (7KO), *Grb10* KO (10KO) and *Grb7*:*Grb10* double knockouts (DKO). Weights are shown for the four offspring genotype groups for (A) Embryo; and (B) Placenta. (C) These values have been used to calculate the embryo to placenta weight ratio as a measure of placental efficiency. Values represent means and SEM, tested by one-way ANOVA using Kruskal-Wallis and Dunn’s post hoc statistical tests. Sample sizes were, for embryo WT n=24, *Grb7* KO n=5, *Grb10* KO n=27, *Grb7*:*Grb10* DKO n=6, and for placenta and embryo:placenta ratio WT n=23, *Grb7* KO n=5, *Grb10* KO n=27, *Grb7:Grb10* DKO n=6. Asterisks indicate *p*-values, *p <0.05, **p <0.01, ****p<0.0001.

Placental weights followed a similar pattern (Figure 7B), with wild type (0.0969±0.0038g) and *Grb7* KO (0.0895g±0.0069g) similar in size to each other, and smaller than *Grb10* KO (0.1119±0.0032g) and *Grb7:Grb10* DKO (0.1165±0.0082g), which were also comparable in size. This meant *Grb10* KO and *Grb7:Grb10* DKO placentae were 15% and 22% larger than wild type, respectively, while *Grb7* KO placentae were 8% smaller. The only statistically significant difference was between wild type and *Grb10* KO samples (p<0.05). Next, the ratio of embryo to placental mass was calculated for each genotype as an estimate of placental efficiency (Figure 7C). Again, the only statistically significant difference from wild type (9.997±0.2546) was an increase in the value for *Grb10* KO (11.3825±0.2775) placental efficiency (p<0.01), which has been observed previously [53], although the value for *Grb7:Grb10* DKO (11.4659±0.7403) was similarly increased. Meanwhile, *Grb7* KO (9.9562±0.7074) placental efficiency was near identical to wild type.

### Loss of *Grb7* has a subtle influence on adult body composition

To characterise the physiology of *Grb7* KO mice we first used dual x-ray absorptiometry (DXA) to compare wild type and KO adults at 15 weeks of age. For both males (Figure 8A-F) and females (Figure 8G-L) there were no obvious differences in any of the measured parameters, including total body weight, lean or fat weights, lean to fat ratio, bone mineral content (BMC) or bone mineral density (BMD). The same animals when physically weighed again showed no differences in total body weight for males (Figure 8M) or females (Figure 8S). A selection of organs was next dissected and weighed, with an emphasis on insulin-responsive tissues. In most cases there were no differences between wild type and *Grb7* KO mice, including in the weights of brain, liver, pancreas, gastrocnemius muscle, tongue and kidneys for both males (Supp. Figure 1A-F) and females (Supp. Figure 1O-T), with the same true for testes in males (Supp. Figure 1G). Unsurprisingly, the same tissues showed no differences between wild type and *Grb7* KO when weights were expressed as a proportion of body weight in males (Supp. Figure 1H-N) or females (Supp. Figure 1U-Z). A second skeletal muscle, the masseter muscle also showed no difference in weight for male (Figure 8N) or female (Figure 8T). The only exceptions to the general rule were two major visceral WAT depots, gonadal and renal, in which the *Grb7* KO weights were lower than wild type for both males (Figure 8O,Q) and females (Figure 8U,W). Reduction of *Grb7* KO fat depot mass in females was significant for renal (Figure 8W; p<0.05) but not gonadal WAT (Figure 8U), although the magnitude of the effect was similar for each depot (*Grb7* KO female depots were 45% and 40% lower than wild type for renal WAT and gonadal WAT, respectively). There was a similar strong trend in males, with the *Grb7* KO gonadal depot 35% smaller and renal 32% smaller than wild type. Relative depot weights were consistent with this pattern, being significant for renal (Figure 8X; p<0.05) but not gonadal (Figure 8V) WAT in females, though not males (Figure 8P,R). The differences in adipose depot weights could not be accounted for by changes in food consumption. Food intake, measured at intervals over 12-14 days, was similar between wild type and *Grb7* KO mice whether expressed simply as daily intake in grams for males (Supp. Figure 2A) and females (Supp. Figure 2D), or as daily food intake adjusted for body weight either at the start of the feeding study period (Supp. Figure 2B,E) or at the end (Supp. Figure 2C,F).

**Figure 8.**
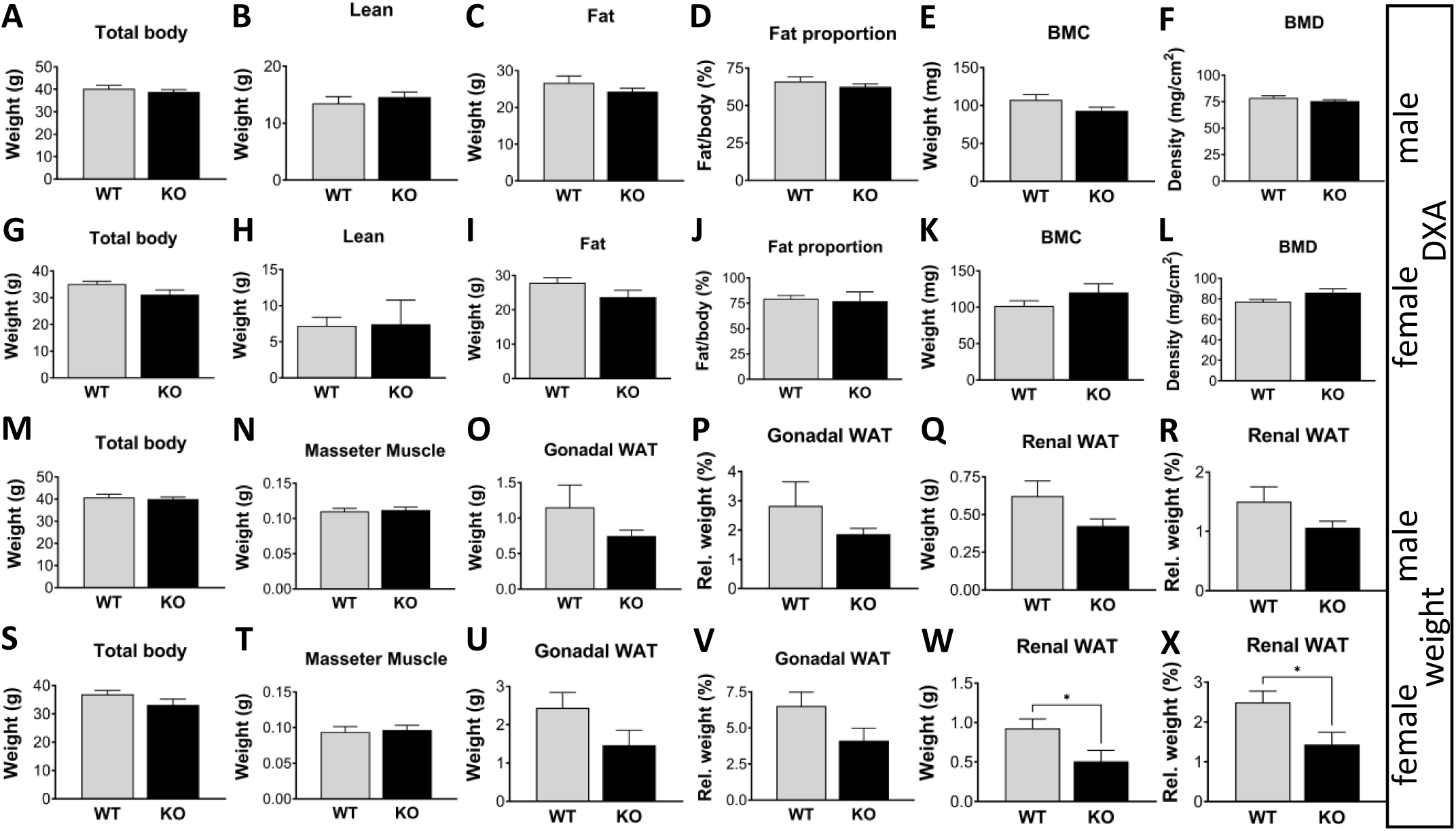
Body composition analysis of adult *Grb7* KO mice. DXA analysis of 15 week old males (A-F) and Females (G-L), showing estimates of total body weight (A, G), lean body content (B, H), fat body content (C, I), fat as a proportion of body weight (D, J), bone mineral content (E, K), and bone mineral density (F, L). For the same animals, physical weights were then obtained for the body and selected tissues and organs for both males (M-R) and females (S-X). Physical weight data are shown for total body (M, S), masseter muscle (N, T); gonadal WAT (O, U) and renal WAT (Q, W), along with weights as a proportion of body weight for gonadal WAT (P, V) and renal WAT (R, X). Graphs show means and SEM, and differences between the means were evaluated using a Student’s t-test. Sample sizes for DXA measurements were, for males WT n=9, *Grb7* KO n=10 and females WT n=5, *Grb7* KO n=3. Sample sizes for physical weight data were, for males WT n=12, *Grb7* KO n=13, and for females WT n=9, *Grb7* KO n=7. Asterisks indicate *p*-values, *p <0.05.

### Sexually dimorphic influence of Grb7 on glucose handling

An initial assessment of glucose handling was carried out by comparing wild type and *Grb7* KO mice at 14 weeks of age. Fasted glucose levels in male *Grb7* KO (7.3±0.5 mM) mice were significantly higher (p<0.05) than wild type (5.7±0.4 mM) (Figure 9A). Levels in free fed animals were near identical in wild type (17.5±1.8 mM) and *Grb7* KO (17.0±2.1 mM) animals (Figure 9B). In a standard glucose tolerance test *Grb7* KO males were slightly slower in clearing a glucose load, however, statistical comparison of the areas under each curve indicated no significant difference (Figure 9C). In contrast, circulating glucose levels in females were similar in both fasted (wild type, 7.8±0.4 mM; *Grb7* KO 7.6±0.6 mM; Figure 9D) and fed (wild type, 16.2±2.3 mM; *Grb7* KO 12.6±4.2 mM) animals (Figure 9E). However, in a glucose tolerance test, *Grb7* KO females cleared a glucose load significantly quicker (p<0.01) than wild type animals as judged by comparison of the areas under each curve (Figure 9F).

**Figure 9.**
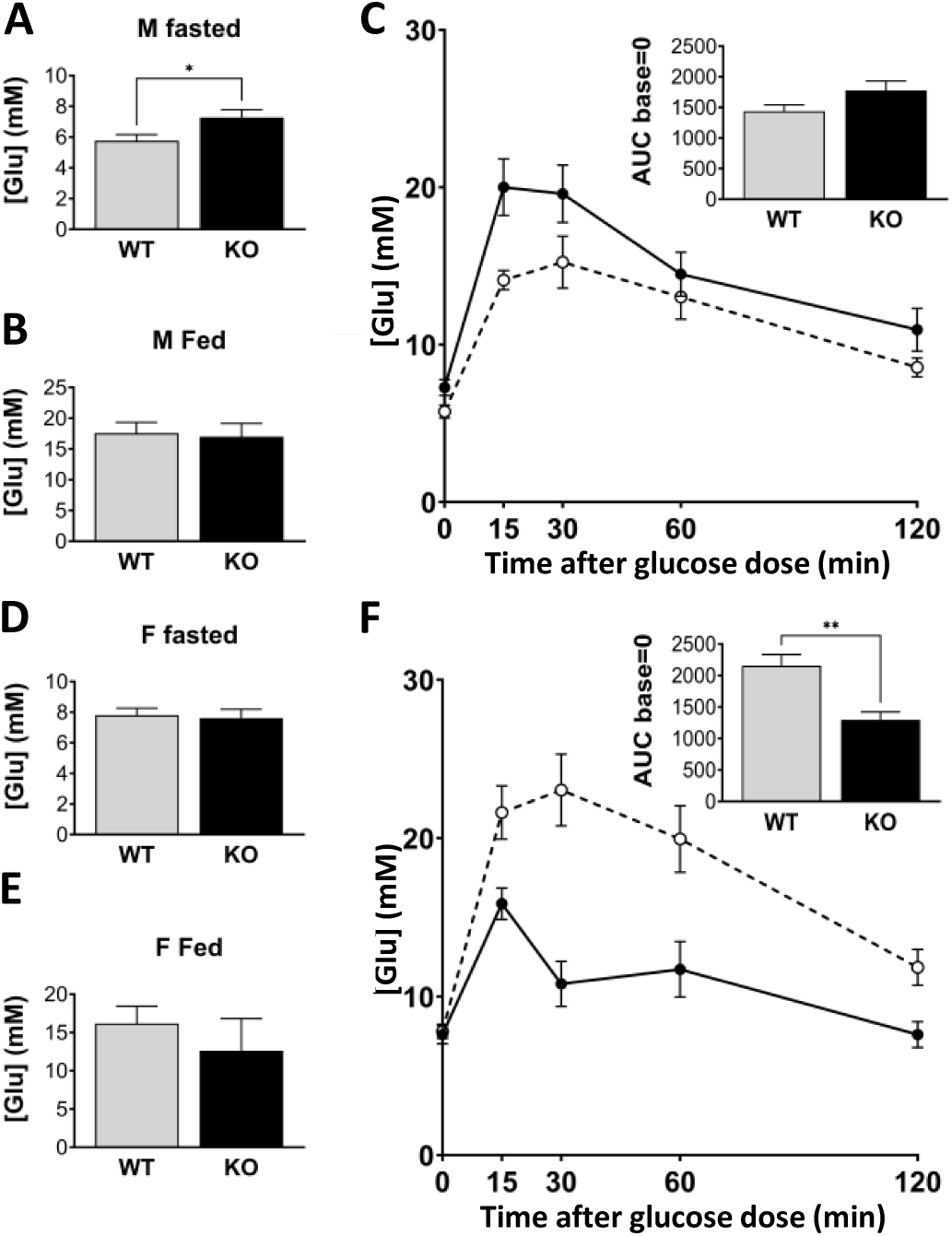
Glucose homeostasis in adult *Grb7* KO mice. Circulating glucose levels were compared for both male (A-C) and female (D-F) wild type and *Grb7* KO mice, aged 14 weeks, in the fasted (A, D) and fed (B, E) states. The same animals were also tested for the ability to clear a glucose load after a fasting period in a standard glucose tolerance test (C, F). Glucose levels were measured at intervals over a time-course of 120 minutes with areas under the curve (AUC) measured in each case for statistical comparison. Graphs show means and SEM, and differences between the means were evaluated using a Student’s t-test. Sample sizes for male fasted and fed glucose levels were WT n=12, *Grb7* KO n=13, for female fasted glucose WT n=9, *Grb7* KO n=7, and fed glucose WT n=9, Grb7 KO n=6. Sample sizes for glucose tolerance tests were, for males WT n=10, *Grb7* KO n=11, and for females WT n=9, *Grb7* KO n=7. Asterisks indicate *p*-values, *p <0.05, **p<0.01.

## Discussion

Available evidence suggests all three Grb7-family members can interact with an overlapping set of tyrosine kinase receptors and other signalling molecules [1, 11]. Physiological functions for Grb10 and Grb14 have previously been established in the regulation of whole body glucose metabolism through direct interaction with the Insr [26, 27, 33, 34]. In addition, the maternal *Grb10* allele exhibits widespread expression in developing mesodermal and endodermal tissues and is an important negative regulator of fetal growth [17, 18]. Here, we tested the idea that either Grb7 or Grb14 might also play a role in fetal growth regulation, either alone or together with Grb10. First, we examined the developmental expression patterns of Grb7 and Grb14 for comparison with each other and that known for Grb10. Antibodies specific to Grb7 and Grb14 provided a snapshot of the distribution of each protein during organogenesis. Our data revealed that Grb7 and Grb14 each had distinct patterns of expression at e14.5 that were more restricted than that of Grb10 [17, 18], but with substantial overlap between the three adaptors. Expression of Grb7 in the developing epithelial structures of the gut, lung and kidney was entirely consistent with a study that examined the spatial distribution of mouse *Grb7* mRNA only in those developing tissues, along with the adult kidney [50]. More broadly, the data are consistent with a spatially resolved expression atlas of mouse development generated using single cell sequencing technology that shows a similar relationship between *Grb7*, *Grb10* and *Grb14* expression patterns throughout organogenesis (e9.5-e16.5)[54]. Grb10 expression also overlaps with Grb7 in pituitary and cartilage and with Grb14 in choroid plexus, skeletal muscle, cardiac muscle and epidermis. In contrast, there was no obvious overlap in expression in the embryo solely between Grb7 and Grb14. This means that our genetic crosses between *Grb10* KO and either *Grb7* KO or *Grb14* KO mice address the majority of potential sites of developmental redundancy. However, there are multiple sites where developmental expression overlaps for all three adaptor proteins, particularly in endodermal organs such as lung, pancreas, liver and the epithelial lining of the digestive tract. Potentially redundant roles of the Grb7-family genes in these tissues merit further investigation, but the generation of *Grb7*:*Grb10*:*Grb14* triple KO animals was beyond the scope of this study.

In addition, we note that in a recent publication (in the press at the time of writing [55]), a different *Grb7* KO strain has been generated that incorporates a *LacZ* insertion at the endogenous locus. This novel strain has been used to describe Grb7 expression in adult tissues but not in the embryo. It is notable that many of the expression features we observed during development were seen in the adult, including the predominantly epithelial expression in organs such as lung, digestive tract and kidney, as well as liver, pancreas and cartilage. The adult expression pattern includes widespread expression in the brain and spinal cord, which we did not see in the embryo, and it will be interesting to establish when neuronal expression of *Grb7* commences. Consistent with our results, Lofgren and Kenny [55] report normal development and survival of *Grb7* KO homozygous mutants. However, offspring of *Grb7* KO females fail to thrive and this correlates with *Grb7* expression in mammary epithelium and a potential defect in function of the lactating gland. This is significant in the context of our earlier finding that *Grb10* is expressed from the maternal allele in lactating mammary epithelium. While offspring of *Grb10* KO (*Grb10^m/+^*) dams survive, they exhibit impaired weight gain, consistent with a mammary gland defect, though we found no obvious morphological defect nor gross changes in milk composition or transfer [56]. Clearly, there is scope for functional redundancy between *Grb7* and *Grb10* in mammary function that deserves further investigation.

Next, we used both *Grb7* KO and *Grb14* KO mouse knockout strains in crosses with a *Grb10* KO strain. The crosses provided an opportunity to determine whether the other two Grb7-family members might play a role in fetal growth regulation, either independently or in a manner redundant with *Grb10*. *Grb10* KO mice lacking expression from the maternal *Grb10* allele are born some 30% larger by weight than wild type littermates [17, 18, 22, 23]. The weight increase is accompanied by greater axial length and skeletal muscle volume, indicative of a general increase in musculoskeletal growth [25]. The fact that different tissues of mesodermal and endodermal origin are overgrown to different extents, such that not all *Grb10* KO organs are proportionate with body size, is consistent with the cell autonomous action expected of an intracellular adaptor protein. The excess growth involves changes in cell proliferation and cell cycle parameters, resulting in *Grb10* KO tissues having more cells rather than larger cells [22, 24] such that, for instance, skeletal muscles at birth contain significantly more myofibers [25]. Here, we observed expression of Grb14 in developing skeletal muscle and cartilage, similar to that previously shown to occur from the maternal *Grb10* allele, which immediately suggested a means for the two factors to act redundantly in fetal growth control. However, analysis of progeny from crosses between *Grb10* KO and *Grb14* KO mice did not support this idea. Total body weight of *Grb10* KO PN1 offspring was 32% heavier than wild type littermate controls, in line with previous studies [17, 18, 22, 23, 56], whereas *Grb14* KO pups were almost indistinguishable from wild type. If the two adaptor proteins were to act redundantly to inhibit fetal growth, the predicted outcome would be for *Grb14*:*Grb10* DKO pups to be larger than *Grb10* KO pups. Instead, *Grb14*:*Grb10* DKO pups were intermediate in size between wild type and *Grb10* KO, without being statistically different to either, providing no evidence for functional redundancy as growth inhibitors. Formally, we cannot rule out the possibility that Grb14 promotes growth only when Grb10 is absent, but this seems unlikely.

Knowing that in *Grb10* KO pups some organs display disproportionate growth, relative to body weight, we examined organs from PN1 progeny to detect any effects of *Grb14* KO on organ growth that might not significantly affect total body weight. In the cross between the *Grb10* KO and *Grb14* KO strains, all organs from *Grb10* KO pups were significantly larger than those from wild type controls, as expected [17, 18, 23]. In each case, *Grb14* KO organs were very similar to wild type, indicating no obvious involvement of Grb14 alone in growth regulation at the level of individual organs. *Grb10*:*Grb14* DKO organs were each intermediate in size between wild type and *Grb10* KO, without being significantly different to either, in almost all cases. The one exception was DKO liver, which was significantly larger than wild type but not statistically different from *Grb10* KO liver.

Importantly, none of the *Grb10*:*Grb14* DKO organs we examined were larger than those from *Grb10* KOs, as would be expected if Grb14 were to act redundantly with Grb10 as a growth inhibitor, allowing us to reject this possibility. As is characteristic of *Grb10* KO pups, both brain and kidney were small relative to total body weight, although both were empirically slightly larger than wild type in this cross. Sparing of the brain can be explained by lack of expression from the maternal *Grb10* allele in the developing CNS, though we note that knockout of the paternal allele also has no significant effect on brain size at PN1 [17, 18, 22]. Brain sparing is a well-known phenomenon, for instance, in the context of poor nutrient availability brain growth can be prioritised at the expense of peripheral tissues in a range of metazoan species from flies to mammals, including humans [57]. By selectively inhibiting growth of peripheral tissues maternal *Grb10* could be an important determinant of brain sparing in mammals. Grb14 expression was largely absent in the e14.5 brain, except in the choroid plexus and leptomeninges, such that any effect it could have on neuronal or glial populations would have to be indirect. Maternal Grb10 expression is also restricted to choroid plexus and leptomeninges within the brain [17, 18], providing scope for functional redundancy with Grb14 in some capacity, but not for a major role in brain growth according to our genetic evidence. In e14.5 kidney, expression of Grb14 was essentially absent, making any role in growth regulation unlikely, either alone or acting redundantly with Grb10.

In contrast to the sparing of brain and kidney, *Grb10* KO neonates display disproportionate overgrowth of the liver, which we have previously associated with excess lipid storage in hepatocytes [17, 18, 22] and to be dependent on Insr signalling [23]. This makes liver an interesting case due to the credentials of both Grb10 and Grb14 as physiological regulators of Insr signalling in adult tissues [26, 27, 33, 34]. However, we saw no evidence for liver enlargement in *Grb14* KO neonates nor exacerbation in *Grb10*:*Grb14* DKO pups of the liver size increase seen in *Grb10* KO single mutants. Despite Grb14 expression being strong in cardiac muscle throughout the heart and in the developing lung epithelia, both organs closely followed the same pattern of weight differences displayed by the whole body. *Grb10* KO heart and lungs were overgrown compared to wild type but in proportion to the increase in body weight, as seen previously [17, 18, 22, 23], *Grb10*:*Grb14* DKO organs were similarly overgrown, while those of *Grb14* KO mice were similar in size to wild type. Overall, the PN1 organ weight data supported the conclusion that *Grb10* alone was responsible for the observed differences in weight between wild types and either single or double mutants in progeny of the cross between the *Grb10* KO and *Grb14* KO strains.

In progeny of the cross between *Grb7* KO and *Grb10* KO mice we again saw the characteristic overgrowth of *Grb10* KO PN1 pups, which had a mean weight 32% heavier than wild type littermates. While *Grb7* KO offspring were very similar to wild type, *Grb7*:*Grb10* DKO offspring were overgrown by 33%, near identical to *Grb10* KOs. The same pattern was observed for e17.5 embryos, where *Grb10* KO and *Grb7*:*Grb10* DKO offspring were again significantly enlarged compared to wild type and *Grb7* KO offspring. The pattern also extended to the placenta, where *Grb10* KO has previously been shown to influence placental size through expansion of the labyrinthine volume, the main site of maternal-fetal exchange [53]. Here, the *Grb10* KO and *Grb7*:*Grb10* DKO placentae were each larger than wild type and *Grb7* KO placentae. These data are consistent with five separate previous crosses, involving two different *Grb10* KO strains, in which the *Grb10* KO embryo and placenta has consistently been enlarged compared to wild type [17, 18, 22, 23]. Collectively the PN1 and e17.5 data indicate *Grb7* has no major role in growth regulation, either alone or in combination with *Grb10*. In contrast to the lack of evidence for an influence of *Grb7* on birth weight, the circulating glucose level of PN1 *Grb7*:*Grb10* DKO pups was significantly lower than the wild type level, suggesting redundancy between *Grb7* and *Grb10* in inhibition of Insr function in the neonate. This contrasted with a lack of such a redundant effect between *Grb10* and *Grb14* in *Grb10*:*Grb14* DKO neonates, in which glucose levels were like those of wild type and either the *Grb10* KO or *Grb14* KO single mutants. This in turn contrasts with evidence that there is an additive effect on glucose regulation in *Grb10*:*Grb14* DKO adults [34].

To check for any organ-specific effects of *Grb7* we examined the weights of selected PN1 organs. In this case there was a clear pattern across the genotypes in which most *Grb10* KO and *Grb7*:*Grb10* DKO organs were similarly enlarged compared to their wild type and *Grb7* KO equivalents. Although statistically indistinguishable, *Grb7* KO organs were consistently slightly smaller than wild type, as was the whole body, raising the possibility that Grb7 may have a subtle positive effect on fetal growth. However, in most cases there was no equivalent difference between *Grb10* KO and *Grb7*:*Grb10* DKO organs, as would be expected when the positive and negative effects of the two genes were combined. Also, the *Grb7* KO organs were empirically smaller than wild type whether *Grb7* expression was seen during development (liver and lungs) or not (brain and heart). The one exception was the kidney, where the reduction in *Grb7* KO weight compared with wild type reached statistical significance and there was a corresponding dip in the weight of *Grb7*:*Grb10* KO relative to *Grb10* KO kidneys. Interestingly, both Grb7 and Grb10 are expressed in the epithelial component of developing kidney, whereas Grb14 is not. Growth of the kidney during nephrogenesis is thought to be driven primarily by expansion of the mesenchyme [58]. Low levels of *Grb10* expression in the metanephric mesenchyme could account for the relatively small increases in kidney weight seen in the crosses presented here and in previous studies [17, 18, 22, 23]. However, the strong expression of both Grb7 and Grb10 in epithelia, together with the results of the cross between *Grb7* KO and *Grb10* KO animals, suggests the two factors influence fetal kidney growth through opposing effects on the developing epithelial structures. In this context it is notable that both Grb7 and Grb10 have been shown capable of interacting with the Ret receptor [59, 60]. The GDNF ligand, expressed in the metanephric mesoderm, interacts with Ret to drive proliferation of ureteric bud tip cells, which is essential for expansion of the branching epithelial structures that make up the collecting duct system for each nephron [61]. In addition, Grb10 has been implicated in NEDD4-mediated E3 ubiquitylation and internalisation of Ret receptors in a human embryonic kidney-derived (HEK293) cell line [62]. This antagonistic effect between Grb7 and Grb10 on kidney growth is an intriguing area for future study.

There was similar overlapping expression of Grb7 and Grb10 in lung epithelia, but without convincing evidence of *Grb7* contributing to lung growth. In this case we cannot rule out redundancy involving both Grb7 and Grb14, since there was Grb14 expression detected in at least a subset of lung epithelial cells. Analogous to the situation in kidney, arborisation of the developing lung epithelium requires signalling from FGF10 in the mesenchyme to FGFR2 in the epithelial bud tips to promote bud outgrowth [63]. All three Grb7 family members have been shown to be capable of binding FGF receptors, with Grb7 and Grb14 having higher affinity than Grb10 in a heterologous *Xenopus laevis* oocyte system [64].

In liver there was again no apparent effect of Grb7 on growth, either in *Grb7* KO or *Grb7*:*Grb10* KO pups despite strong Grb7 expression throughout the e14.5 liver suggesting, as for *Grb14*, a lack of *Grb7* involvement in the Insr-mediated accumulation of excess lipid seen in the disproportionately enlarged *Grb10* KO neonatal liver [22, 23]. However, since Grb14 expression was also seen in developing hepatocytes, it remains possible that Grb7 and Grb14 could influence hepatic growth, either through expansion of cell number or lipid storage, in a redundant manner. Further, liver was a prominent site of expression for all three Grb7-family adaptor proteins. In at least two mouse models *Grb10* expression has been linked with hepatic steatosis [65, 66], a precursor of non-alcoholic fatty liver disease (NAFLD) and the most frequent cause of liver failure worldwide [67]. Liver-specific knockout of *Grb10* is sufficient to prevent hepatic lipid accumulation induced by diet or tunicamycin treatment [66]. Similarly, adenoviral-mediated knockdown of mouse *Grb14* in liver resulted in enhanced hepatic Insr signalling while dramatically reducing lipogenesis [68]. A recent genome-wide association study linked both *GRB14* and *INSR* with NAFLD [69], raising the possibility that Grb14 (and potentially all three Grb7-family proteins) could also control lipogenesis in hepatocytes through their ability to regulate Insr signalling.

In the case of pancreas, we have shown that all three adaptor proteins are expressed in the adult organ, specifically in the endocrine cells of the islets of Langerhans, as well as in the developing organ. Expression of Grb10 in pancreatic islets has been previously reported (e.g. [26]) and islet-specific *Grb10* KO mice were found to have increased beta cell mass associated with increased beta cell proliferation, enhanced Insr signalling in islets and elevated insulin secretion, with associated improvement in whole body glucose clearance [29, 70]. In keeping with this, at least one genome wide association study has identified human *GRB10* as a major determinant of endocrine pancreas function [71]. *GRB7* has been linked with several tissues in the context of cancer progression, including liver (hepatocellular carcinoma [72]) and pancreas [73], and a contribution to Insr signalling functions in these tissues cannot be excluded. The strong Grb7 expression we observed in the developing gut epithelium is interesting in view of evidence that *GRB7* has an oncogenic role in cancers of the intestinal tract, including those of the oesophagus [74], stomach [75] and colon [76].

The *Grb7* KO strain, generated specifically for this study, was designed to be a null allele. Consistent with this, we showed that Grb7 was absent in adult tissues normally expressing readily detectable levels of the protein. Our study includes an initial evaluation of adult tissue and organ proportions, as well as whole body glucose handling, in these *Grb7* KO mice. Both male and female *Grb7* KO animals were indistinguishable from wild type littermates as judged by DXA measurements for body mass, lean and fat mass, plus bone mineral content and density. This was largely corroborated by physical weight measurements for a range of tissues and organs. We included two different skeletal muscles (masseter and gastrocnemius) and the muscular tongue because of increased lean mass and muscle weights seen in *Grb10* KO mice [22, 25–27, 34], finding no differences between wild type and *Grb7* KO tissues. Similarly, there were no differences in the weights of brain, liver, pancreas or testes. In contrast, in *Grb7* KO females the renal WAT depot was significantly lighter than that from wild type littermates and the gonadal WAT followed the same trend. In males, both WAT depots were lighter than wild type, but not significantly so. Overall, there was a tendency for *Grb7* KO mice at 15 weeks of age to have reduced visceral adipose despite there being no discernible change in food intake. An explanation for the reduced WAT mass could be a tendency towards increased energy expenditure. Involvement of *Grb10* in the regulation of thermogenesis by brown adipose tissue (BAT) has been shown following adipose-specific *Grb10* KO [77]. In these mice, which lack *Grb10* expression in BAT and WAT, WAT depots become enlarged due to mTORC1-dependent reduction of lipolysis in WAT and reduced thermogenic activity in BAT. The more convincing adipose deficit in *Grb7* KO females was accompanied by a significant improvement in glucose clearance rate, whereas *Grb7* KO males tended to clear glucose less efficiently than wild types, if anything, and had significantly higher fasted glucose levels, unlike females. The reasons for these sexual dimorphisms are unclear, but it is interesting to note that *GRB14* has been linked with adiposity traits specific to women in genome wide association studies designed to identify sexually dimorphic traits [78, 79]. Both male and female *Grb10* KO mice exhibit modest improvements in glucose and insulin sensitivity, despite reduced circulating insulin levels [22, 26]. Studies of glucose metabolism in *Grb14* KO have so far been restricted to male mice, which have improved glucose clearance, again with reduced insulin levels [33, 34]. *Grb10*:*Grb14* DKO males were resistant to the impairment in glucose tolerance induced by a high fat diet whereas the *Grb10* KO and *Grb14* KO single mutants were not [34]. Considering all these findings, it will be interesting to establish whether the difference in adipose mass becomes more prominent as *Grb7* KO animals age and to examine body composition and glucose handling in adult compound mutants, combining *Grb7* KO with *Grb10* KO and *Grb14* KO.

In conclusion, we can reject the idea that *Grb7* or *Grb14* act as inhibitors of fetal growth, either alone or in combination with *Grb10*, and instead *Grb7* may positively influence fetal kidney growth, acting antagonistically to *Grb10*. In contrast, we have provided evidence that *Grb7* can contribute to the regulation of whole-body glucose handling along with *Grb10* and *Grb14*. This places all three signalling adaptor proteins in the frame as important regulators of energy homeostasis and potential therapeutic targets for major metabolic disorders including obesity, type 2 diabetes and NAFLD.

## Declarations

### Ethical approvals

Experiments involving mice were subject to local ethical review by the University of Bath Animal Welfare and Ethics Review Board and carried out under licence from the United Kingdom Home Office. The manuscript has been written as closely as possible in accordance with the Animal Research: Reporting of In Vivo Experiments (ARRIVE) guidelines (https://arriveguidelines.org/).

## Consent for publication

Not applicable.

## Availability of data and materials

The datasets used and analysed during the current study are available from the corresponding author on reasonable request.

## Competing interests

The authors declare that they have no competing interests.

## Funding

This work was supported by Medical Research Council grants [MR/S00002X/1, MR/S008233/1]. The funder had no specific role in the conceptualization, design, data collection, analysis, decision to publish, or preparation of the manuscript.

## Author contributions and rights retention

AW conceived the project, collected and interpreted the data, wrote the manuscript and assembled the figures. FMS and ASG set up genetic crosses, collected data and contributed to data analysis. KM carried out the statistical analyses and generated the final graphs and images. RJD and LJH contributed to manuscript revision. All authors read and approved the final manuscript. For the purpose of open access, the author has applied a Creative Commons Attribution (CC BY) licence to any Author Accepted Manuscript version arising from this submission.

## Acknowledgements

For generously supplying the *Grb7* KO mouse strain we thank Benjamin Margolis (University of Michigan, USA). We are grateful to University of Bath Biological Services Unit staff for outstanding animal care.

## Figure Legends

**Supplementary Figure 1.**
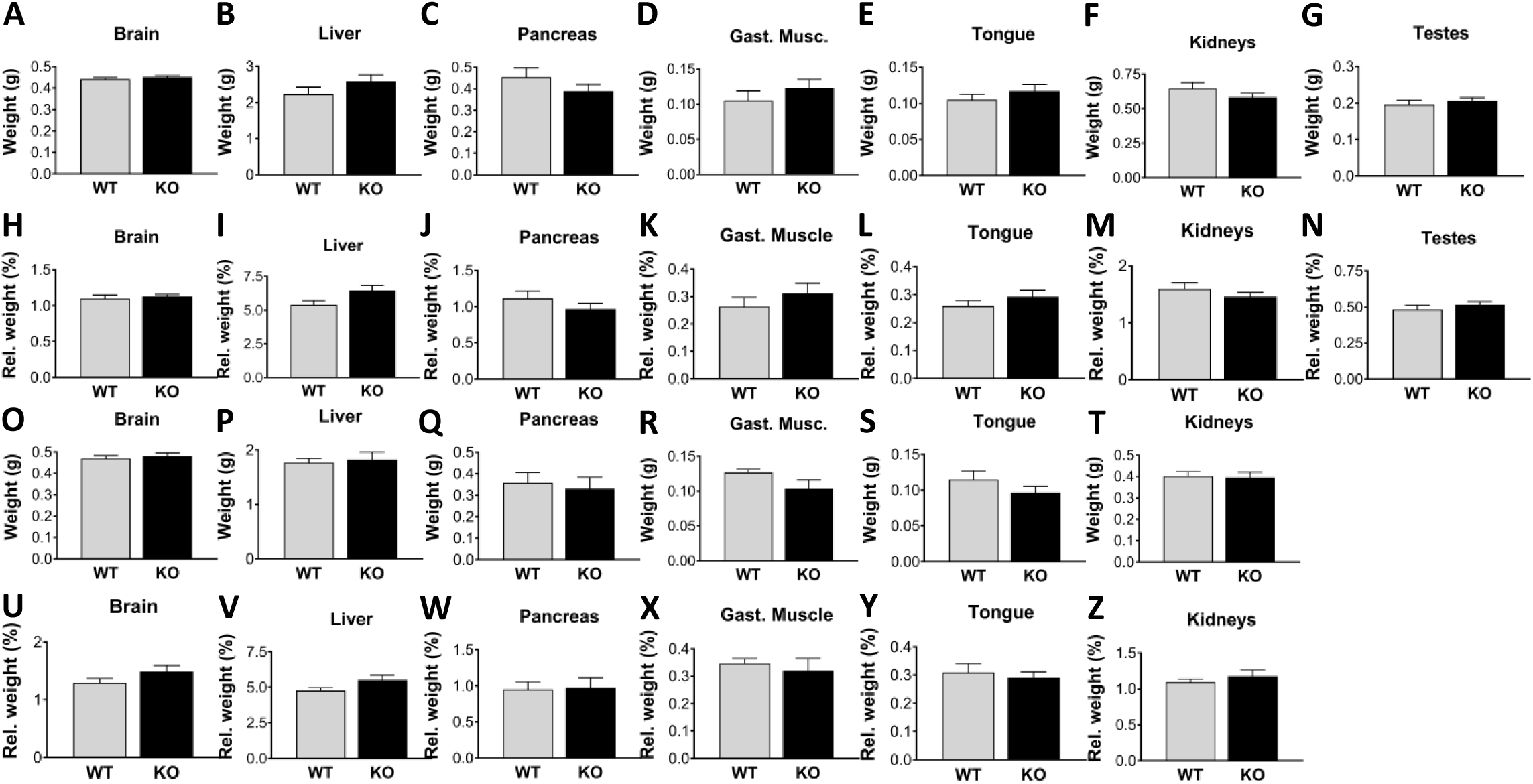
Organ and tissue weight data for adult *Grb7* KO mice. Additional tissue and organ weights were obtained for the same animals described in Figure 8. Raw weights are shown for both males (A-G) and females (O-T), alongside the weight of each tissue as a proportion of body weight for males (H-N) and females (U-Z). The selected tissues and organs were, brain (males A,H; females O,U), liver (males B,I; females P,V), pancreas (males C,J; females Q,W), gastrocnemius muscle (males D,K; females R, X), tongue (males E,L; females S,Y), kidneys (males F,M; females T,Z) and testes (males G,N). Graphs show means and SEM, and differences between the means were evaluated using a Student’s t-test except where variance differed between the WT and *Grb7* KO in which case a Mann Whitney test was used (Male gastrocnemius muscle, gonadal WAT, renal WAT, and relative brain, gastrocnemius muscle, gonadal WAT and renal WAT.). For males, sample sizes were WT n=12, *Grb7* KO n=13, except for testes WT n=11, *Grb7* KO n=13. For females, sample sizes were WT n=9, *Grb7* KO n=7, except for kidneys WT n=9, *Grb7* KO n=6. In all cases differences between wild type and *Grb7* KO were non-significant (p>0.05).

**Supplementary Figure 2.**
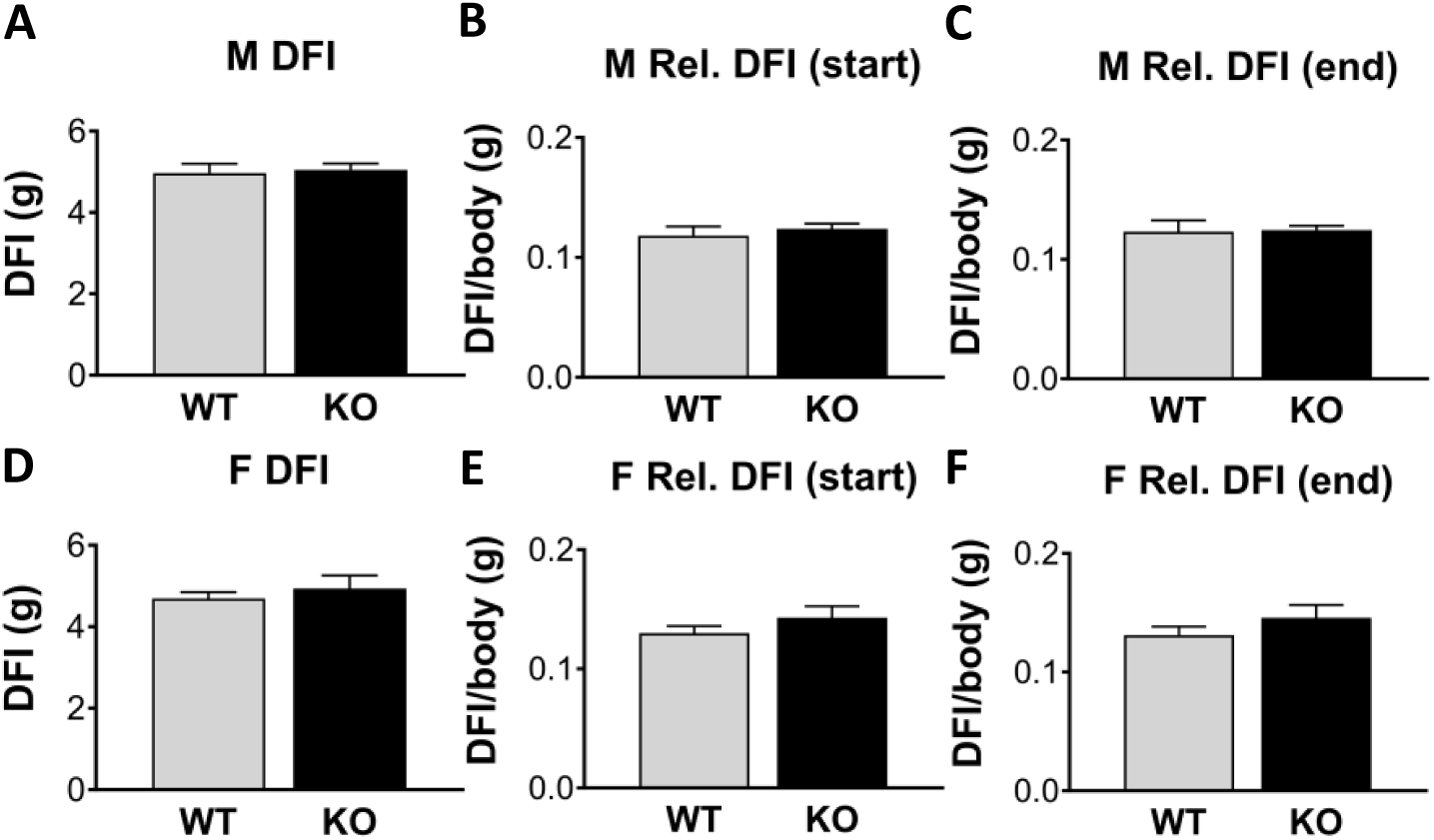
Daily food intake (DFI) measurements for adult *Grb7* KO mice. The mean weight of food consumed per day by animals of 12-14 weeks of age is shown for males (A) and females (D). Food consumption was also expressed as a proportion of body weight, both at the start of the experiment, for males (B) and females (E), and at the end of the experiment, for males (C); and females (F). Graphs show means and SEM, and differences between the means were evaluated using a Student’s t-test except in the case of Male DFI (end), which was analysed using a Mann Whitney test because the variance of WT and *Grb7* KO groups differed significantly. Sample sizes were, for males WT n=12, *Grb7* KO n=13, and for females WT n=9, *Grb7* KO n=7. In all cases differences between wild type and *Grb7* KO were non-significant (p>0.05).

**Supplementary Table 1.**
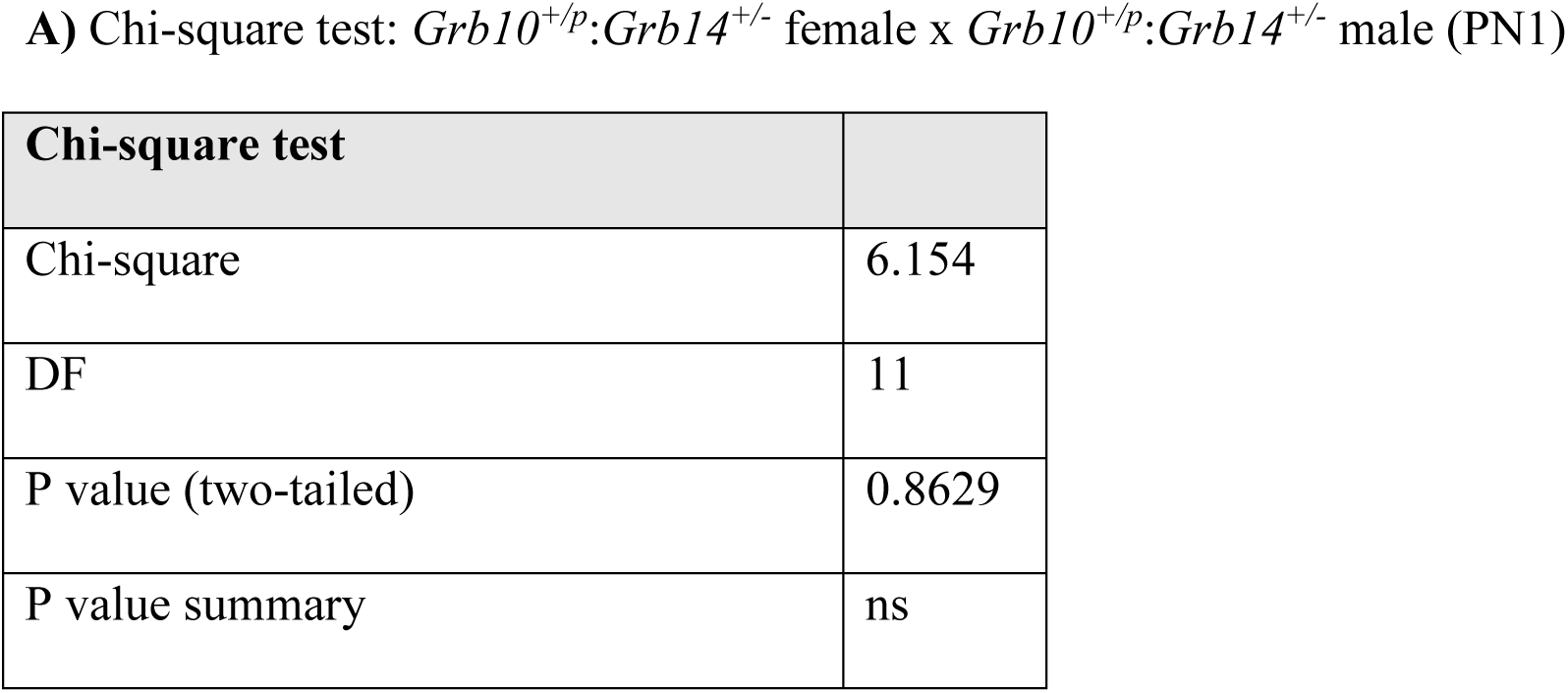
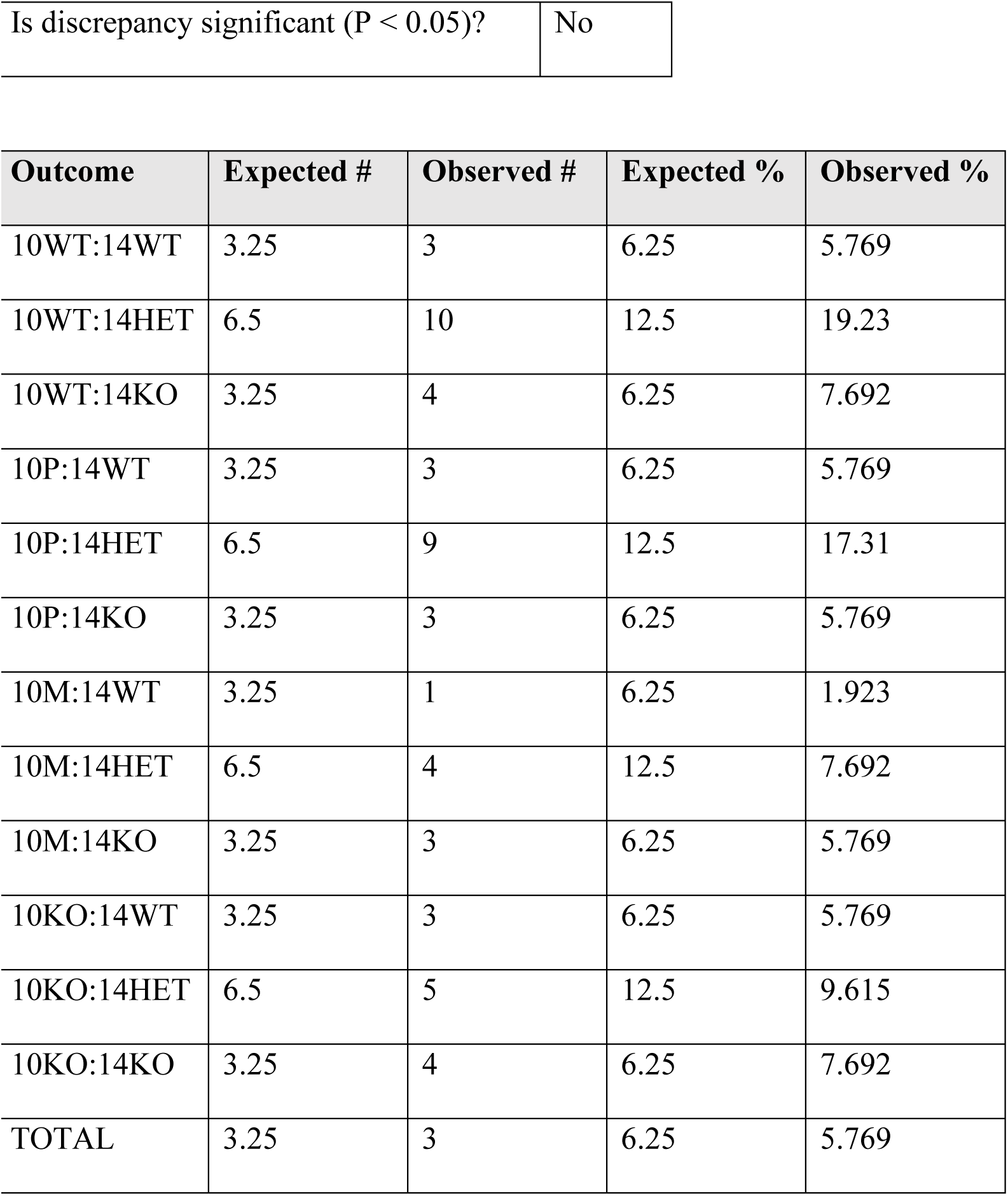
Chi-squared statistical tests of offspring survival from crosses between *Grb10* KO and *Grb14* KO mouse strains. Offspring collected from crosses between *Grb10^+/p^*:*Grb14^+/−^* females x *Grb10^+/+^*:*Grb14^+/−^* males at PN1. Deviation from the expected Mendelian ratio was considered significant at p<0.05.

**Supplementary Table 2.**
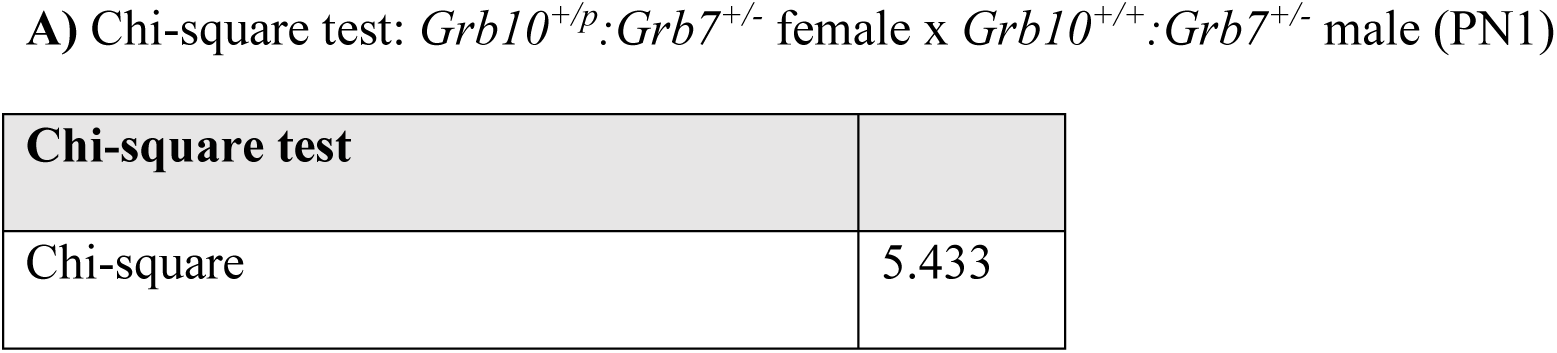
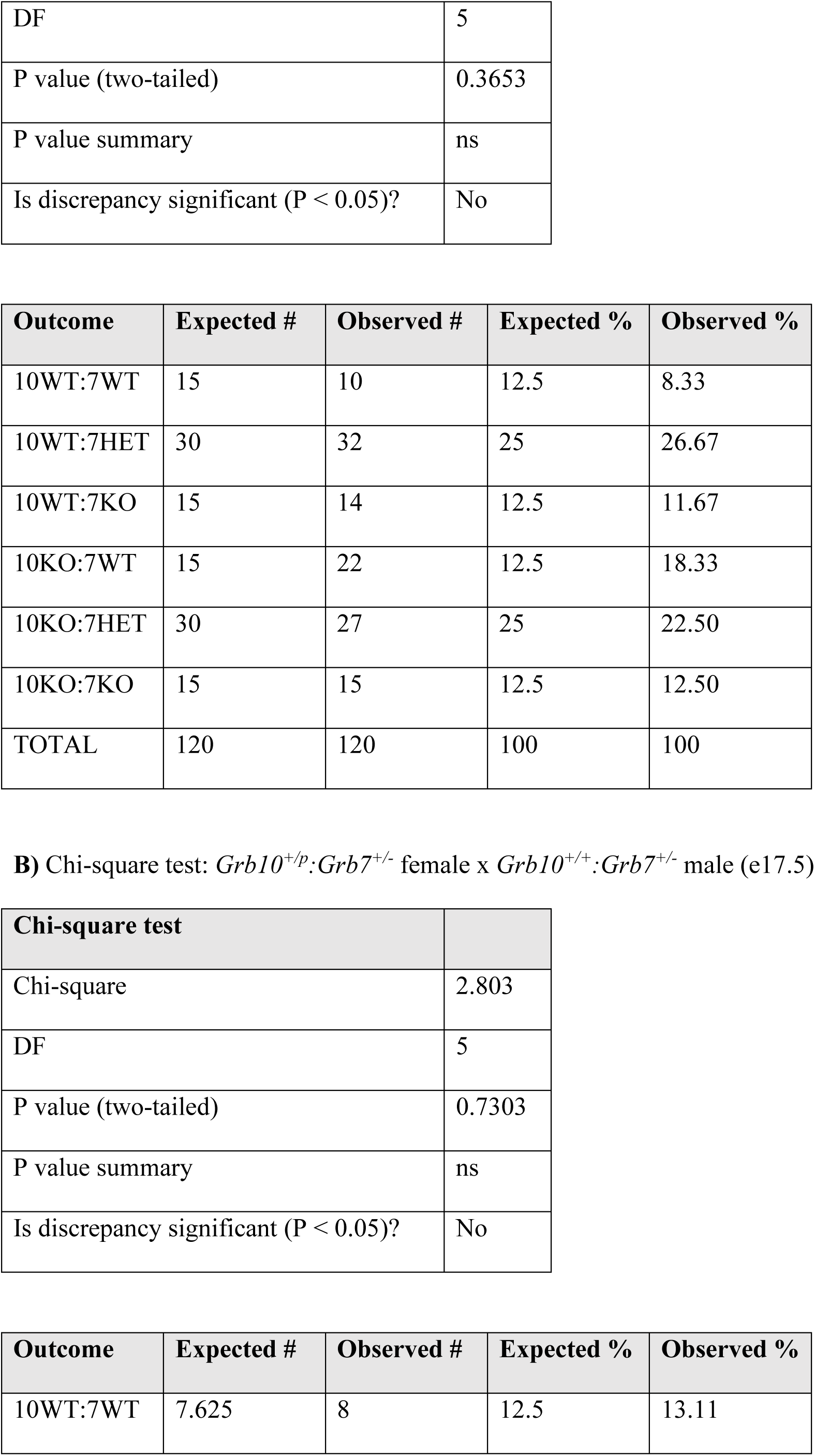
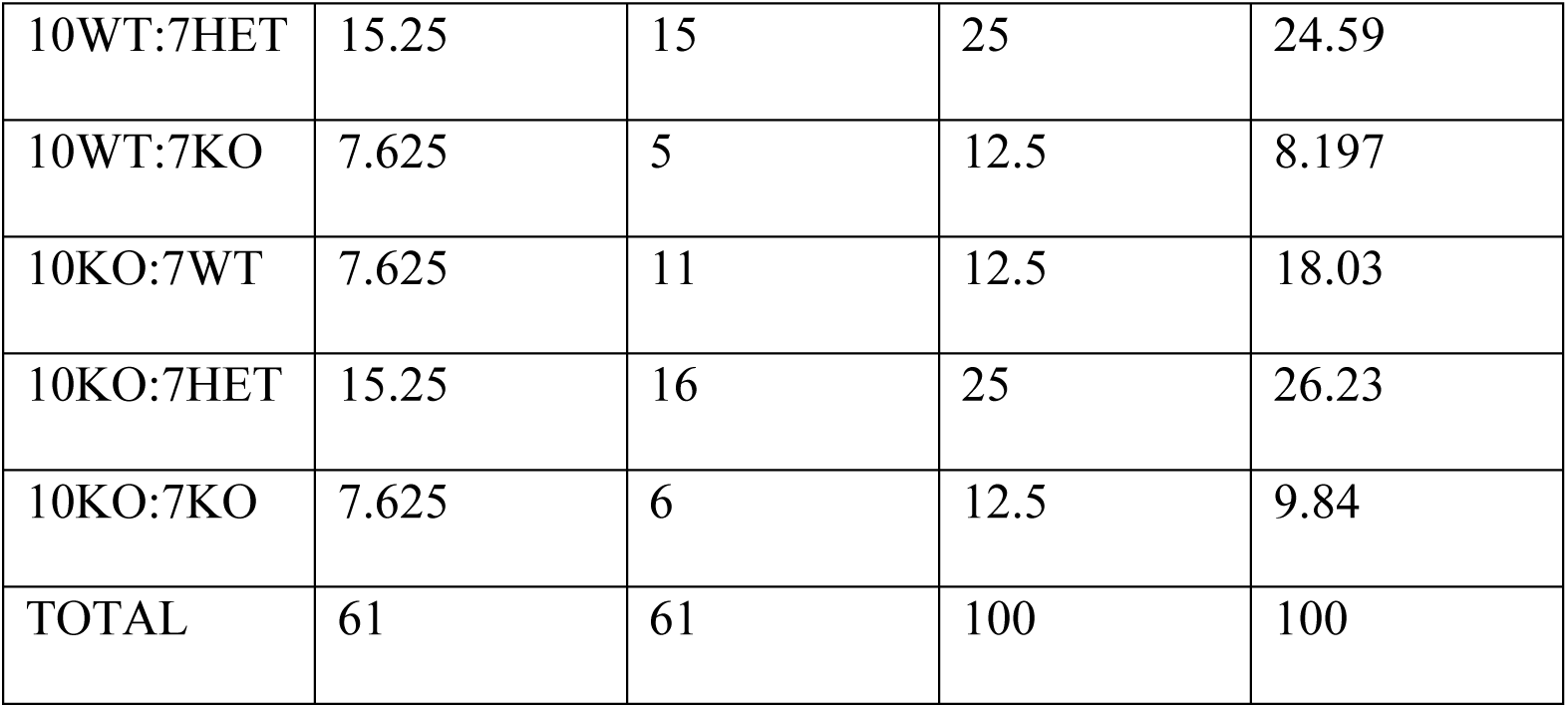
Chi-squared statistical tests of offspring survival from crosses between the *Grb10* KO and *Grb7* KO strains. Offspring collected from crosses between *Grb10^+/p^:Grb7^+/−^* females x *Grb10^+/+^:Grb7^+/−^*males at PN1 (A) and at e17.5 (B). Deviation from the expected Mendelian ratio was considered significant at p<0.05.

## Notes

### Competing Interest Statement

The authors have declared no competing interest.

