## Supplemental Tables for "*Grb7*, *Grb10* and *Grb14,* encoding the growth factor receptor-bound 7 family of signalling adaptor proteins have overlapping functions in the regulation of fetal growth and post-natal glucose metabolism"

**Supplementary Tables**

| **Chi-square test** | |
|---|---|
| Chi-square | 6.154 |
| DF | 11 |
| P value (two-tailed) | 0.8629 |
| P value summary | ns |
| Is discrepancy significant (P < 0.05)? | No |

| **Outcome** | **Expected #** | **Observed #** | **Expected %** | **Observed %** |
|---|---|---|---|---|
| 10WT:14WT | 3.25 | 3 | 6.25 | 5.769 |
| 10WT:14HET | 6.5 | 10 | 12.5 | 19.23 |
| 10WT:14KO | 3.25 | 4 | 6.25 | 7.692 |
| 10P:14WT | 3.25 | 3 | 6.25 | 5.769 |
| 10P:14HET | 6.5 | 9 | 12.5 | 17.31 |
| 10P:14KO | 3.25 | 3 | 6.25 | 5.769 |
| 10M:14WT | 3.25 | 1 | 6.25 | 1.923 |
| 10M:14HET | 6.5 | 4 | 12.5 | 7.692 |
| 10M:14KO | 3.25 | 3 | 6.25 | 5.769 |
| 10KO:14WT | 3.25 | 3 | 6.25 | 5.769 |
| 10KO:14HET | 6.5 | 5 | 12.5 | 9.615 |
| 10KO:14KO | 3.25 | 4 | 6.25 | 7.692 |
| TOTAL | 3.25 | 3 | 6.25 | 5.769 |

**Supplementary Table 2.** Chi-squared statistical tests of offspring survival from crosses between the *Grb10* KO and *Grb7* KO strains. Offspring collected from crosses between

*Grb10*$^{+/p}$*:Grb7*$^{+/-}$ females x *Grb10*$^{+/+}$*:Grb7*$^{+/-}$ males at PN1 (A) and at e17.5 (B). Deviation from the expected Mendelian ratio was considered significant at $p<0.05$.

**A)** Chi-square test: *Grb10*$^{+/p}$*:Grb7*$^{+/-}$ female x *Grb10*$^{+/+}$*:Grb7*$^{+/-}$ male (PN1)

| **Chi-square test** | |
|---|---|
| Chi-square | 5.433 |
| DF | 5 |
| P value (two-tailed) | 0.3653 |
| P value summary | ns |
| Is discrepancy significant (P < 0.05)? | No |

| **Outcome** | **Expected #** | **Observed #** | **Expected %** | **Observed %** |
|---|---|---|---|---|
| 10WT:7WT | 15 | 10 | 12.5 | 8.33 |
| 10WT:7HET | 30 | 32 | 25 | 26.67 |
| 10WT:7KO | 15 | 14 | 12.5 | 11.67 |
| 10KO:7WT | 15 | 22 | 12.5 | 18.33 |
| 10KO:7HET | 30 | 27 | 25 | 22.50 |
| 10KO:7KO | 15 | 15 | 12.5 | 12.50 |
| TOTAL | 120 | 120 | 100 | 100 |

**B)** Chi-square test: *Grb10*$^{+/p}$*:Grb7*$^{+/-}$ female x *Grb10*$^{+/+}$*:Grb7*$^{+/-}$ male (e17.5)

| **Chi-square test** | |
|---|---|
| Chi-square | 2.803 |
| DF | 5 |
| P value (two-tailed) | 0.7303 |

| | |
|---|---|
| P value summary | ns |
| Is discrepancy significant ($P < 0.05$)? | No |

| **Outcome** | **Expected #** | **Observed #** | **Expected %** | **Observed %** |
|---|---|---|---|---|
| 10WT:7WT | 7.625 | 8 | 12.5 | 13.11 |
| 10WT:7HET | 15.25 | 15 | 25 | 24.59 |
| 10WT:7KO | 7.625 | 5 | 12.5 | 8.197 |
| 10KO:7WT | 7.625 | 11 | 12.5 | 18.03 |
| 10KO:7HET | 15.25 | 16 | 25 | 26.23 |
| 10KO:7KO | 7.625 | 6 | 12.5 | 9.84 |
| TOTAL | 61 | 61 | 100 | 100 |
